# An Inordinate Fondness for Species with Intermediate Dispersal Abilities

**DOI:** 10.1101/644872

**Authors:** Ben Ashby, Allison K. Shaw, Hanna Kokko

**Author notes:** Authors contributed equally to the manuscript (Ben Ashby, Allison K. Shaw).

## Abstract

JBS Haldane is widely quoted to have quipped that the Creator, if one exists, has an inordinate fondness for beetles. Although Coleoptera may not be the most speciose order once Hymenopteran diversity is fully accounted for, as a whole the very clear differences in species diversity among taxa require an explanation. Here we show both analytically and through stochastic simulations that dispersal has eco-evolutionary effects that predict taxa to become particularly species-rich when dispersal is neither too low nor too high. Our models combine recent advances in understanding coexistence in niche space with previously verbally expressed ideas, where too low dispersal imposes biogeographic constraints that prevent a lineage from finding new areas to colonize (reducing opportunities for speciation), while too high dispersal impedes population divergence, leading to few but widely distributed species. We show that this logic holds for species richness and is robust to a variety of model assumptions, but peak diversification rate is instead predicted to increase with dispersal. Our work unifies findings of increasing and decreasing effects of dispersal rate on speciation, and explains why organisms with moderate dispersal abilities have the best prospects for high global species richness.

## Introduction

Dispersal can influence biodiversity through both ecological and evolutionary processes. Ecologically, dispersal can enhance colonization, which increases local diversity if all else is equal (MacArthur and Wilson 1967). Yet all else is typically not equal: increased movement, by increasing contact with similar species, can limit diversity through competitive exclusion (Macarthur and Levins 1967). When multiple processes interact, the outcome can be scale-dependent: theory suggests that regional diversity should decrease with increasing dispersal (Hubbell 2001, Mouquet and Loreau 2003), while local diversity is often maximized at intermediate dispersal (unimodal pattern) (Mouquet and Loreau 2002, 2003). In some cases this relationship can also become monotonic or multimodal (Haegeman and Loreau 2014). Experimental studies that manipulate dispersal have typically found a unimodal relationship between dispersal and diversity maintenance, such that diversity peaks at intermediate dispersal (Kneitel and Miller 2003, Cadotte 2006, Venail et al. 2008).

Evolutionarily, dispersal can influence the speciation process itself, and perhaps, may provide an explanation as to why certain taxa (as surmised in Haldane’s quip about beetles) are more speciose than others. Increased dispersal ability can increase the extent of species ranges (Jablonski 1986), allow individuals to colonize new areas, and thus increase opportunities for speciation to occur. However, by increasing population mixing, dispersal can also erode local adaptation and prevent speciation (Mayr 1963). Empirical patterns relating dispersal to speciation rates (or species richness, often interpreted as evidence of speciation rates) are mixed (Coyne and Orr 2004). Marine invertebrates typically show higher speciation rates associated with lower dispersal (Jablonski 1986, Palumbi 1992), while avian studies have found speciation rates to peak at high (Owens et al. 1999, Cockburn 2003), low (Belliure et al. 2000, Claramunt et al. 2012, Weeks and Claramunt 2014), or intermediate dispersal (‘short-distance colonists’ *sensu* Diamond et al. 1976). In angiosperms, speciation rate is related to the dispersal mechanism: lineages with biotic dispersal (e.g. bird-dispersed fruits) tend to have higher diversity than those that rely on abiotic methods such as wind (Ricklefs and Renner 1994, Dodd et al. 1999, Price and Wagner 2004). This pattern may partly reflect biotic dispersers achieving higher dispersal distances, but the interpretation is complex because biotic dispersers can also have an improved likelihood of arriving in a suitable location.

The evolutionary role of dispersal in speciation has received much less theoretical attention than the ecological role of dispersal in biodiversity maintenance. In broad agreement with Mayr’s ideas (Mayr 1963), recent theoretical work has shown that increased dispersal rate can, by promoting gene flow, increase the time to a first speciation event (Gavrilets et al. 2000), and reduce speciation rates for species that, as a result of good dispersal ability, occupy large geographic areas (Birand et al. 2012). These unidirectional predictions conflict with a number of recent empirical studies, which instead employ a verbal model arguing that speciation rates, and in turn species richness, should be highest at intermediate dispersal distances (Price and Wagner 2004, Claramunt et al. 2012, Agnarsson et al. 2014, Schenk and Steppan 2018) (Fig. 1). The core of this idea is expressed succinctly by Price and Wagner (Price and Wagner 2004): ‘Species-rich lineages may have moderate dispersability that is effective enough to extend the geographic range of whole lineages, yet infrequent enough to depress levels of gene flow’ (see also Vermeij 1987, Bleiweiss 1990). Especially if habitable environments are patchy (e.g. Hawaiian islands for angiosperms, Price and Wagner 2004), too little movement will lead to much of the world remaining undiscovered by many of the potential lineages that could persist there, while too much movement maintains gene flow at a level that constrains speciation. Diversity remains low for different reasons at either end: it remains limited in the first case as species stay put in narrow-range sympatry, and in the latter case low diversity exists in a pattern of few species each occupying a very large range.

**Figure 1.**
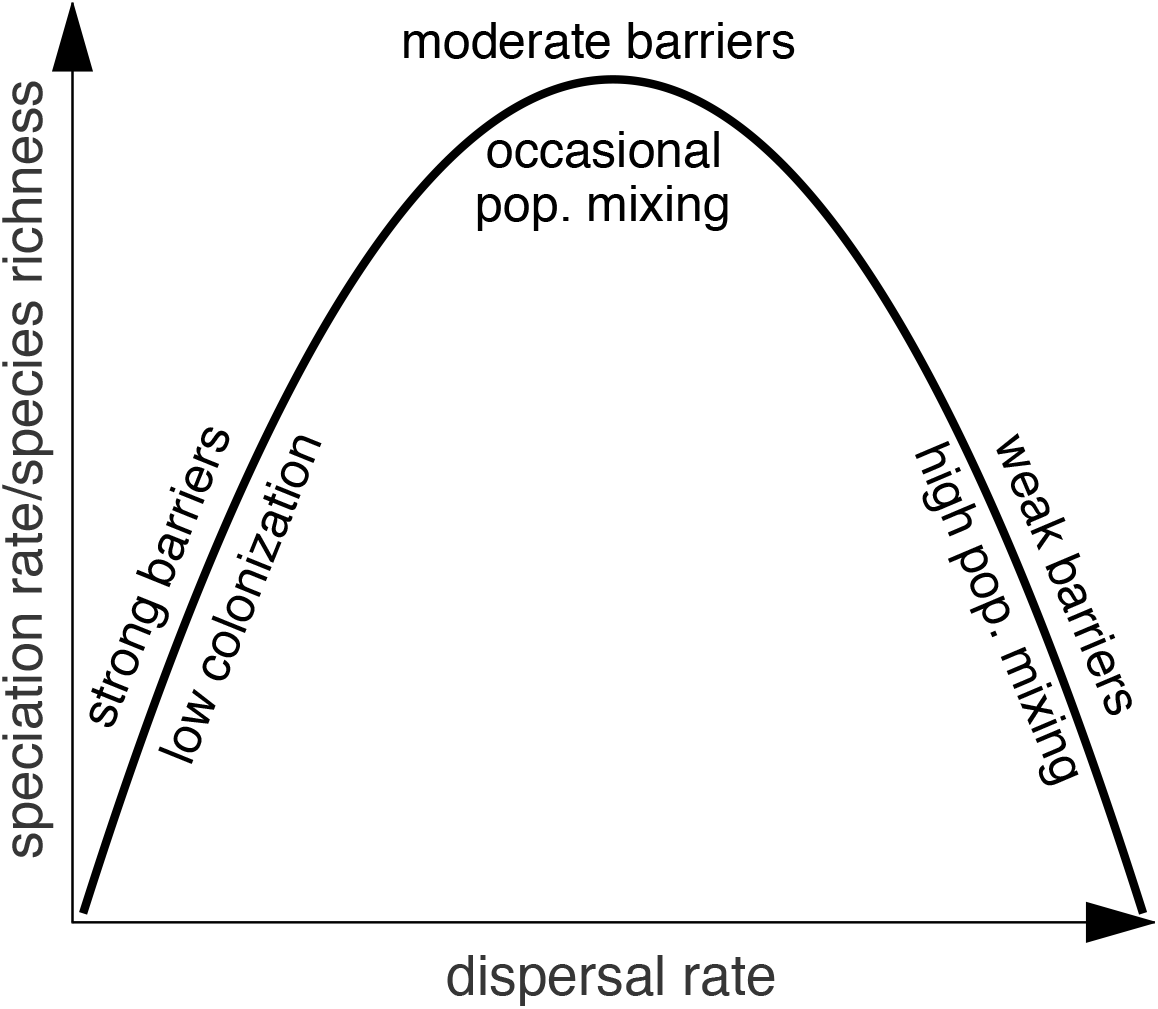
The ‘intermediate dispersal model’ of speciation/species richness. When dispersal is rare colonization is too low and when dispersal is high the metapopulation behaves similarly to a single population. Speciation rates and species richness are therefore predicted to peak at intermediate dispersal rates (Price and Wagner 2004, Claramunt et al. 2012, Agnarsson et al. 2014).

Here we investigate if and when intermediate dispersal leads to the best prospects for speciation, as suggested by some of the verbal models, or whether the negative effect of dispersal on speciation is bound to dominate the outcome, as suggested by the two theoretical contributions mentioned above (Gavrilets et al. 2000, Birand et al. 2012). We use a combination of deterministic and stochastic metapopulation models to explore the relationship between dispersal and speciation, showing: (1) analytically that species richness peaks at intermediate dispersal when dispersal increases colonization and the correlation between patches, and (2) how the nature of resource competition interacts with dispersal to influence the peak diversification rate, species richness, and the proportion of the potential niche space that is filled. Finally, we return to Haldane’s quip about beetles (Farrell 1998), speculating that our results hint at potential causality, and discuss insights from other taxonomic groups.

## Material and Methods

### Deterministic toy model

To illustrate the argument, we first propose a toy model of dispersal and speciation with a large number of possible species and patches, which tracks the proportion of patches that are occupied, *p*(*t*), and the average proportion of species within occupied patches that are unique, *u*(*t*) (i.e. given a set of species distributed across multiple patches, we can calculate the proportion of species within each occupied patch that are not found in any other patches and then take the average value). The toy model is given by:

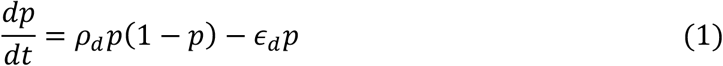

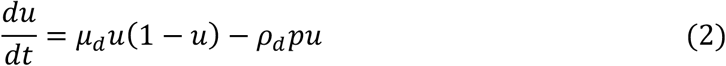

where *ρ*_*d*_ is the dispersal rate per occupied patch, *∊*_*d*_ is the patch extinction rate (i.e. the rate at which patches become unoccupied due to environmental catastrophes) and *μ*_*d*_ is the speciation rate per patch. This model’s take on dispersal and speciation is simplistic, but it mathematically captures the essential features of the verbal model: (1) patch colonisation increases with dispersal; and (2) dispersal reduces the differences in species distributions between patches. Since the model is continuous, the product of *p*(*t*) and *u*(*t*) gives a measure of the diversity of species analogous to species richness, which is maximised when all patches are occupied (*p*(*t*) = 1) and no species are found in more than one patch (*u*(*t*) = 1).

This toy model provides a very simplistic view of dispersal and speciation: it lacks important features of real populations such as resource competition, it views dispersal and speciation in a very simplified manner, and it does not predict speciation rates. Furthermore, it views extinction as a patch-level process, whereby if species do not disperse at a sufficient rate then they will eventually be driven extinct due to a localized catastrophe. These assumptions suffice for a basic check of the verbal model, but to gain a deeper understanding of the relationship between dispersal and species diversification we explore a more detailed stochastic metapopulation model with dispersal between a finite number of patches, local resource competition, and species-rather than patch-level extinctions.

### Stochastic metapopulation model

We adapt the multidimensional niche model described by Ashby et al. (Ashby et al. 2017) to allow for (1) multiple patches with dispersal and (2) neutral traits to distinguish between species that evolve independently in different patches. A total of *L* patches are arranged in a ring (to remove boundary effects) with dispersal occurring between adjacent patches (Fig. 2A). Individuals compete locally over *y* ‘substitutable’ or ‘non-substitutable’ resource types (*sensu* Ashby et al. 2017). It is assumed that organisms can replace one type of substitutable resource with another (e.g. different food sources), but cannot do so with resources that are non-substitutable (e.g. food and nest sites). In practice this means that the competition kernels (see below, equations 3–4) are determined multiplicatively for substitutable resources and additively for non-substitutable resources (Fig. 2B-D). As a result, differentiation along one niche axis may be sufficient to allow coexistence when resources are substitutable, but differentiation along all axes is required resources are non-substitutable. Indeed, previous modelling of niche evolution (without dispersal) has shown that the distinction between the two resource types is crucial, with substitutable resources leading to a densely-packed niche space with overlapping species distributions (Fig. 2E), whereas non-substitutable resources lead to a sparsely-packed niche space with non-overlapping species distributions across each niche axis (Fig. 2F) (Ashby et al. 2017).

**Figure 2.**
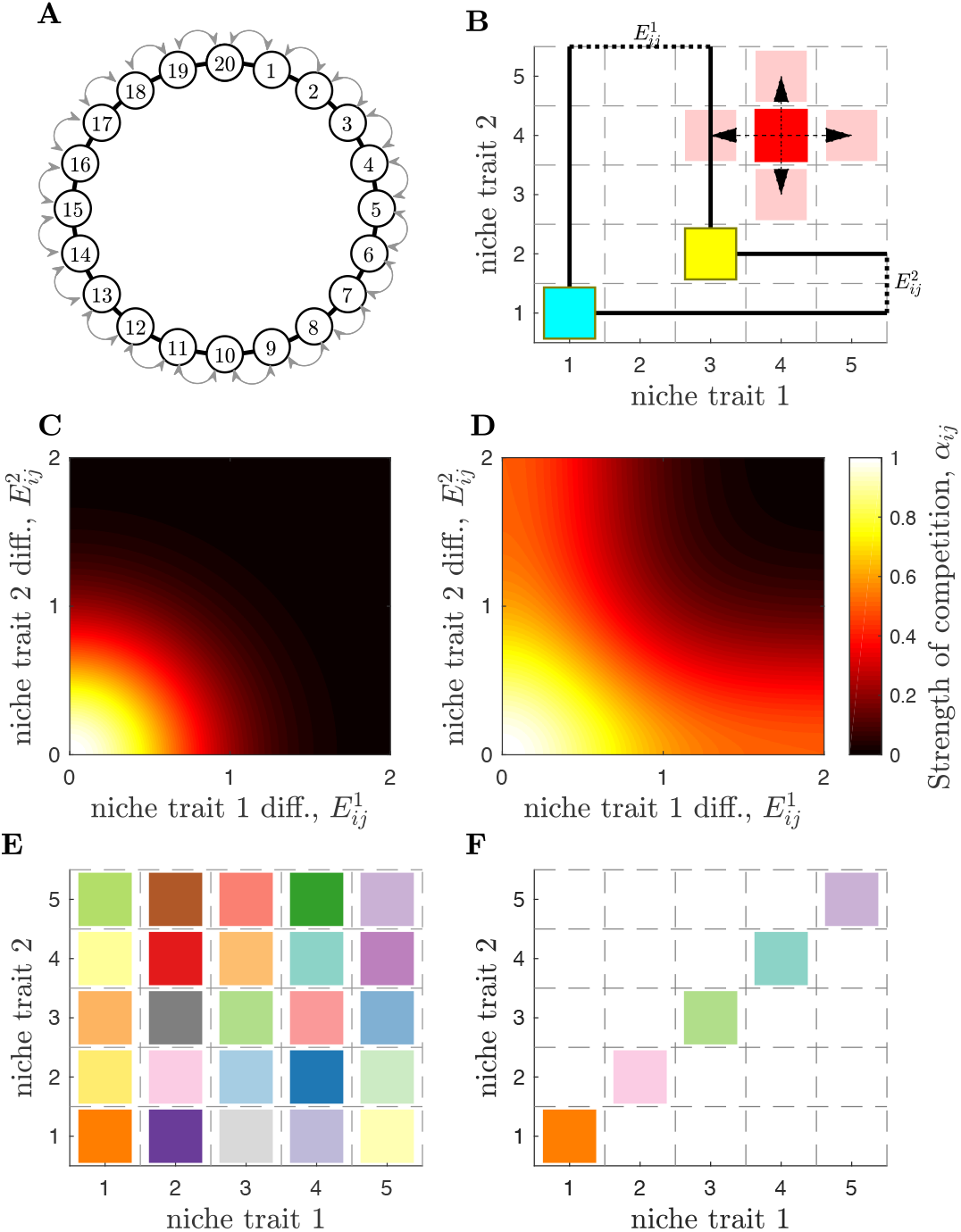
Stochastic metapopulation model overview. (A) The environment is split into *L* = 20 patches arranged in a ring to remove boundary effects. Competition occurs within patches and dispersal between adjacent patches (arrows). (B) Each species occupies a position in the niche space corresponding to its preferred resources/traits. Speciation (red) occurs at rate *μ*_*s*_ causing a change in one niche dimension (arrows). The strength of competition, *α*_*ij*_, between two species (yellow and blue) is based on the distance between species in each dimension of the niche space, 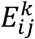, and whether the resources/traits are: (C) ‘substitutable’ (e.g. different food sources), or (D) ‘non-substitutable’ (e.g. food source and nest sites), as described in the main text (*sensu* Ashby et al. 2017). (E) When resources are substitutable, species self-organise to occupy all potential niches. (F) In contrast, species must differentiate across non-substitutable resource axes, producing non-overlapping distributions with many potential niches unfilled.

We assume each resource type is subdivided into *c*_*k*_ resources (*k* = 1, …, *y*) giving a maximum of 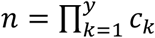 potential niches, which may be filled or unfilled by organisms in each patch (*sensu* Ashby et al. 2017). In other words, each niche is defined by a unique set of niche traits which a species may potentially possess, regardless of whether a species currently exists with those traits. We define 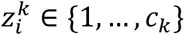 to be the preferred resource within resource type *k* for species *i* (each resource type is also arranged in a ring to remove boundary effects). Species are defined by their set of niche traits, 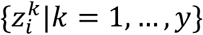, combined with a neutral trait, *v*_*i*_. In our preliminary simulations we allowed the neutral trait to mutate at a given rate, but we found this to be more computationally intensive and to produce qualitatively similar results to a simpler method. We instead set the neutral trait for each species to correspond to the patch in which the species first arose, *v*_*i*_ ∈ {1, …, *L*}, giving a maximum of *s* = *Ln* potential species. We include a neutral trait so that two lineages in different patches which independently evolve to occupy the same niche are classed as distinct species (detectable as their neutral traits having different values).

We define 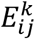 to be the distance between the preferred resources of species *i* and *j* within resource type *k* (Fig. 2B) and the strength of competition between species *i* and *j* to be:

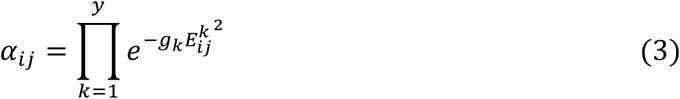

for substitutable resources (Fig. 2C) and

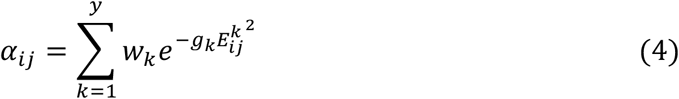

for non-substitutable resources (Fig. 2D), where *g*_*k*_ > 0 mediates the niche breadth for resource type *k* (i.e. the extent to which species consume resources similar to their preferred resource) and *w*_*k*_ is the relative importance of the resource type when resources are non-substitutable 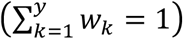.

The number of individuals belonging to species *i* in patch *p* at time step *t* is given by 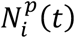. Each time step is assumed to encompass many generations implying a timescale over which species distributions change due to niche overlap. The population size change between time steps (each step is equal to one time unit) due to ecological processes (births, deaths, resource competition) is given by

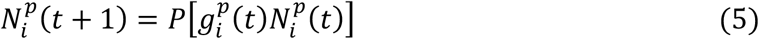

where *P*[*λ*] is a Poisson distributed random variable with mean *λ*. The growth function, 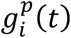, is the expected proportional change in population size due to competition with other organisms in the same patch and is equal to:

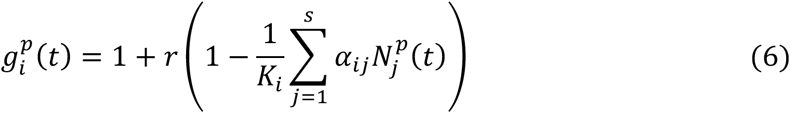

where *r* scales the rate of the within-patch dynamics, *α*_*ij*_ ∈ [0,1] is the degree of niche overlap (i.e. strength of competition) between species *i* and *j*, and *K*_*i*_ is the baseline carrying capacity of species *i*, which is drawn from a normal distribution with mean *K*_*mean*_ and standard deviation *K*_*std*_.

Once the within-patch dynamics have been updated we allow speciation, dispersal, and random extinctions to occur. The total number of speciation events from species *j* to *i* is

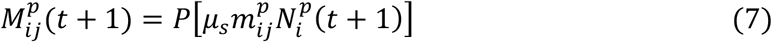

where *μ*_*s*_ scales the speciation rate and 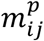 determines if species *j* can mutate to species *i*: 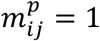 if the species are exactly one niche trait apart from each other 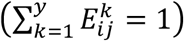 and if *v*_*i*_ = *p* so that the new species’ neutral trait corresponds to the patch in which it arises (to distinguish it from other species that evolve in other patches to occupy the same niche). Since species are identified by a unique set of traits, speciation is modelled as a single mutation in one of these traits (this could be considered a model of asexual lineages arising through mutation), and so the patch-level speciation rate is proportional to the local population size (i.e. mutation supply). The number of dispersal events from patch *p* to patch *u* for species *i* is given by:

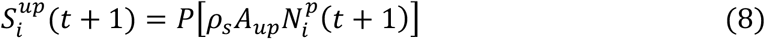

where *ρ*_*s*_ is the dispersal rate and *A*_*up*_ is the adjacency matrix, such that *A*_*up*_ = 1 if patches *u* and *p* are adjacent and *A*_*up*_ = 0 otherwise. The total change in species *i* in patch *p* due to dispersal is therefore:

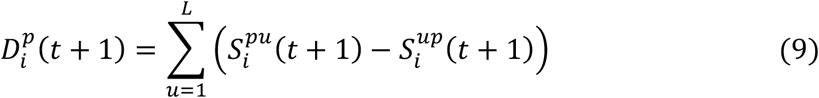

Finally, we allow random extinction events to occur with probability *∊*_*s*_ per species per patch at each time step.

We focus our analysis on the dispersal rate (*ρ*_*s*_) and the type of resource competition (substitutable or non-substitutable resources), carrying out 1,000 simulations per parameter set (source code available in the *Supplementary material*). We initially seed one patch with a single species, then simulate the dynamics for *T* = 10^4^ time steps. The duration was chosen based on preliminary simulations, which typically reached quasi-equilibrium within this time period (longer simulations do not qualitatively change the results, as shown in the *Supporting Information*).

We measure species richness at time *t*, *R*(*t*), by pooling all patches together and counting species that are above a threshold of *δ* = 10 individuals. We calculate the quasi-equilibrium species richness, *R*\*, by taking the mean of *R*(*t*) over the final 1,000 time steps of the simulation. To help understand the process that leads to changes in species richness, we calculate the peak diversification rate, *σ*, by sampling the population every 100 time steps and counting the difference in the total number of extant species across the entire metapopulation between sampling points. This allows us to determine the conditions that lead to rapid increases in species richness. Finally, to understand the relationship between species richness and resource competition, we calculate the proportion of potential niches that are filled at the quasi-equilibrium. This allows us to determine how species patterns are driven by the total number of distinct niches discovered by the globally diversifying taxon.

## Results

### Deterministic toy model

At the non-trivial equilibrium, we have 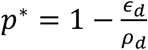 and 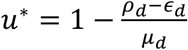 for *ρ*_*d*_ > *∊*_*d*_ and *μ*_*d*_ > *ρ*_*d*_ − *∊*_*d*_, with species diversity across the patches equal to the product of *p*\* and *u*\*. Given that the proportion of occupied patches (*p*\*) increases with dispersal (*ρ*_*d*_) and the average proportion of species that are unique within each occupied patch (*u*\*) decreases with dispersal, species diversity is maximised at intermediate dispersal (Fig. 3), showing analytically that the core principles of the verbal model are mathematically sound (Fig. 1).

**Figure 3.**
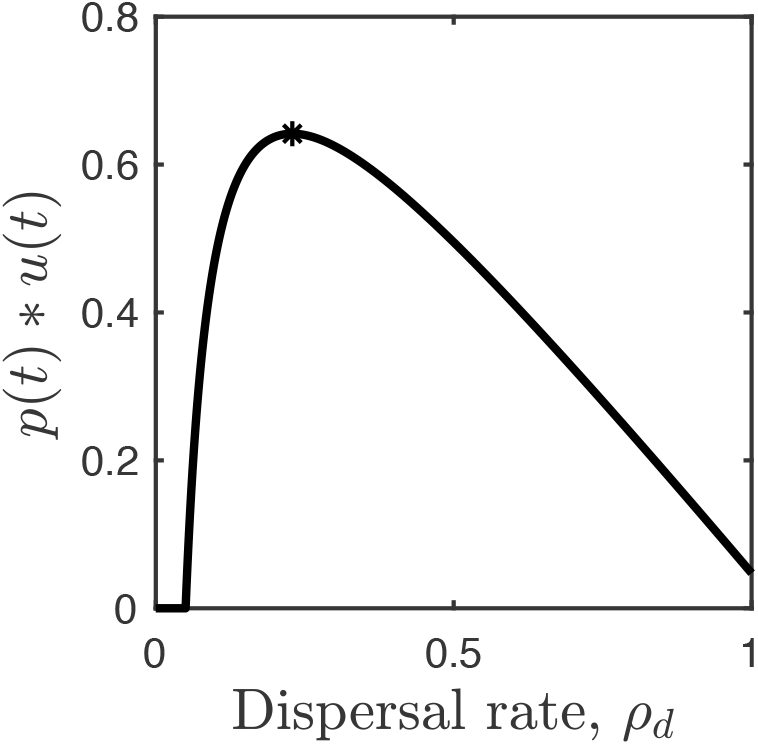
Results for the deterministic toy model. The diversity of species, given by the product of the proportion of patches that are occupied, *p*(*t*), and the average number of species within each occupied patch that are unique to that patch, *u*(*t*), peaks at intermediate dispersal (star): 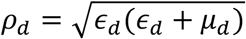. Illustrative parameters: *∊*_*d*_ = 0.05, *μ*_*d*_ = 1.

### Stochastic metapopulation model

We separate our analysis of the stochastic metapopulation model into two parts, considering the effects of dispersal first on the quasi-equilibrium (QE) species richness, *R*\*, and niche space, and then on the maximum speciation rate, *σ*, that underlies and partially explains the emerging species richness. In line with both the verbal and deterministic models, we find that species richness is typically maximised at intermediate dispersal (Fig. 4A; parameter values specified in figure captions), a finding that is extraordinarily robust across different modes of resource competition (i.e. substitutable or non-substitutable resources), a wide range of model parameter values (Fig. S1–S7), and over longer time scales (Fig. S8).

**Figure 4.**
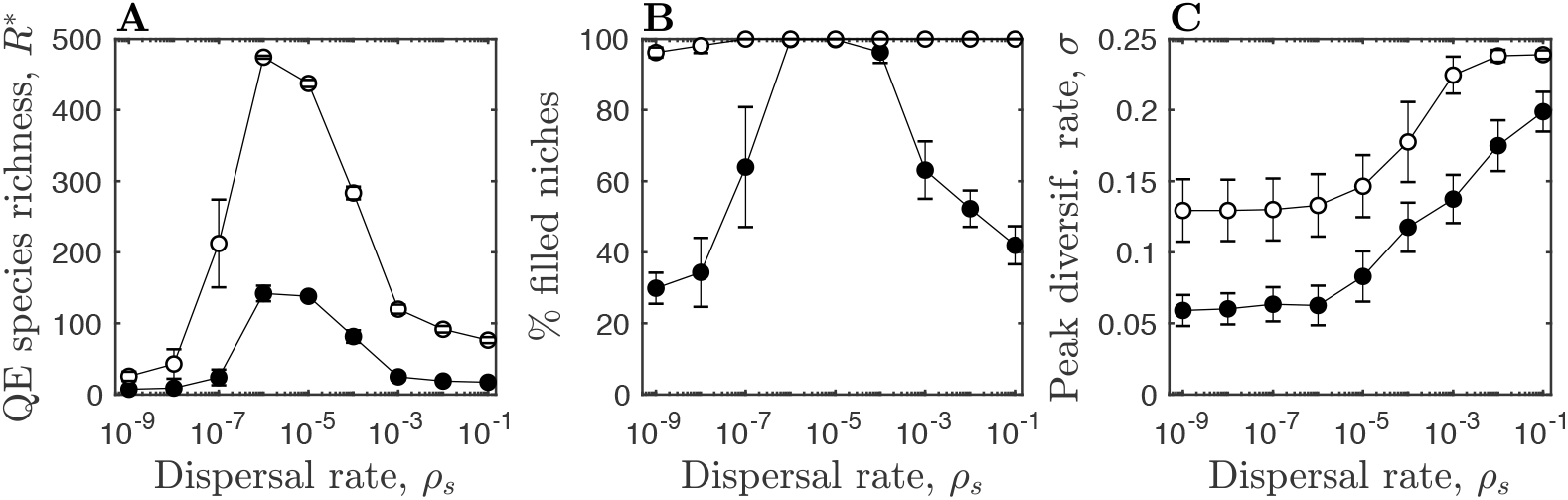
Simulation results. Panels show results for substitutable (white) and non-substitutable resources (black) as a function of the dispersal rate, *ρ*_*s*_. Markers correspond to mean values from 1,000 simulations, with error bars showing one standard deviation. (A) Quasi-equilibrium (QE) species richness, *R*\*; (B) percentage of potential niches that are filled in at least one patch. (C) Peak diversification rate, *σ*. Parameters: *g*_1_, *g*_2_ = 1.5, *k*_*mean*_ = 1000, *K*_*std*_ = 10, *L* = 20, *c*_1_, *c*_2_ = 5, *n* = 2, *w*_1_, *w*_2_ = 1/2, *∊*_*s*_ = 10^−3^, *μ*_*s*_ = 10^−4^.

The pattern for species richness is qualitatively similar for substitutable and non-substitutable resources, but we find that the extent to which potential niches are filled at the global level depends both on dispersal and resource type. Specifically, the proportion of potential niches that are filled peaks at intermediate dispersal when resources are non-substitutable (e.g. food source and nest sites), but is either roughly constant or increasing when resources are substitutable (e.g. different food sources; Fig. 4B, S1–S8). The reason for this difference lies in the way species are distributed within each patch (Fig. 5). Competition for substitutable resources tends to lead to most niches being occupied within each patch, thus dispersal is not a strong predictor of how much of niche space is filled globally. In some cases, within-patch niche diversity may remain lower when dispersal is rare (due to extinctions that are not compensated for by frequent colonization, Fig. S1B–S8B), while all other dispersal values simply allow maximal niche diversity to be reached (Fig. 4B).

**Figure 5.**
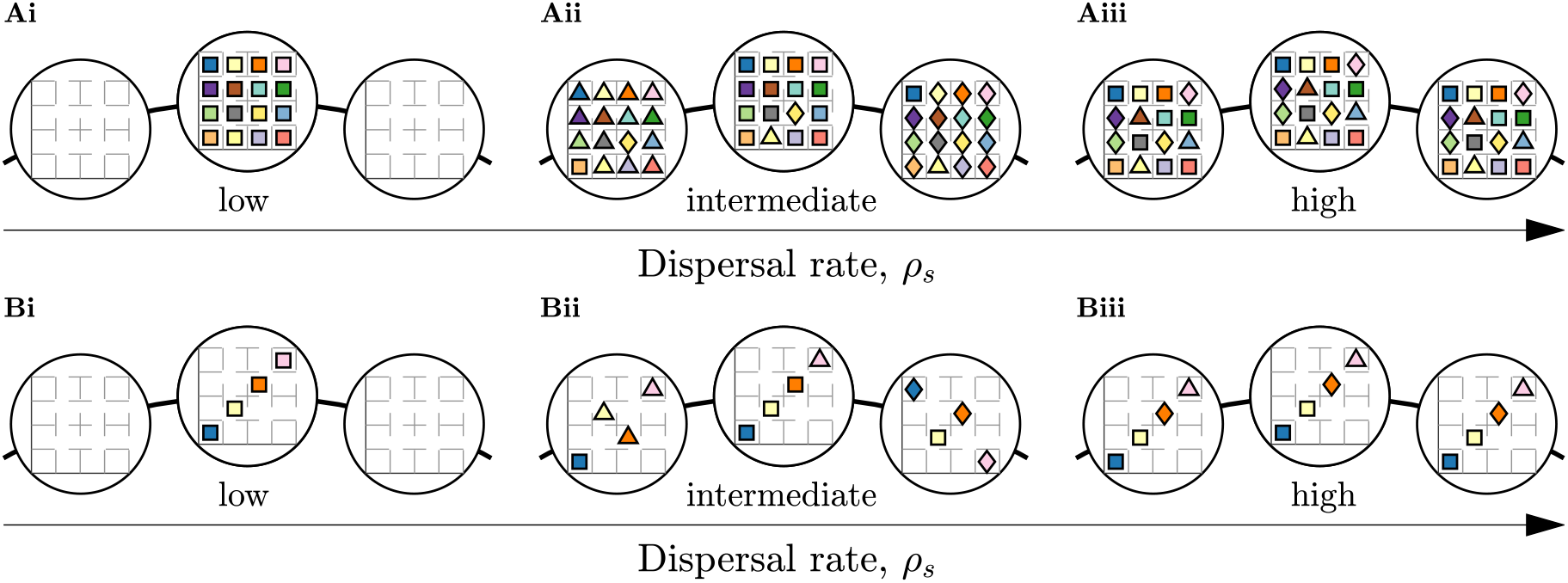
Example interactions between resource type and dispersal at a quasi-equilibrium (end of a simulation). Patches are represented by large connected circles, and markers represent species in a 2-dimensional niche space, with colors corresponding to the niche and shapes to the neutral trait of that species. (A) When resources are substitutable (e.g. different food sources), all potential niches are filled within each patch and so dispersal does not affect the global distribution of occupied niches. (B) When resources are non-substitutable (e.g. food source and nest sites), species separate into non-overlapping distributions within each patch. However, due to the arbitrary nature of the non-overlapping distributions, different species distributions can evolve in each patch at intermediate dispersal, leading to a corresponding peak in the proportion of potential niches that are filled at the global level.

In contrast, competition for non-substitutable resources leads to non-overlapping and potentially arbitrary associations between resources in different patches as an emergent property of the system (Fig. 5). At low or high dispersal, most of niche space remains unfilled since few patches are occupied or there is high mixing between patches eroding variation in niche diversity, respectively. At intermediate dispersal, however, many patches are occupied and the occupied niches between patches are unlikely to be the same due to the arbitrary nature of the non-overlapping distributions. This leads to a peak in the proportion of potential niches that are occupied. In summary, competition for substitutable resources leads to complete saturation of all potential niches whenever a patch is occupied, with no differences between occupied patches, whereas competition for non-substitutable resources leads to arbitrary non-overlapping niches filled within each patch and a peak in occupied niches at the global level for intermediate dispersal.

While the verbal model is well-supported in terms of species richness (and in terms of occupied niches when resources are non-substitutable), we find that the peak diversification rate increases with dispersal (Fig. 4C). This is because the peak diversification rate depends on both the total number of species across the system and the local availability of new niches (Fig. 6). When dispersal is low, the peak diversification rate is highly constrained: few patches are occupied, thus few new species can arise and persist due to the limited number of extant species and available niches. As dispersal increases, more patches become colonized and so there are more niches available locally, causing the peak diversification rate to increase. When dispersal is high, all patches rapidly become occupied, and so the number of available niches increases very quickly, leading to a high peak diversification rate. In fact, the explosion of diversity can initially outpace local resource competition before less-fit species are gradually driven extinct.

**Figure 6.**
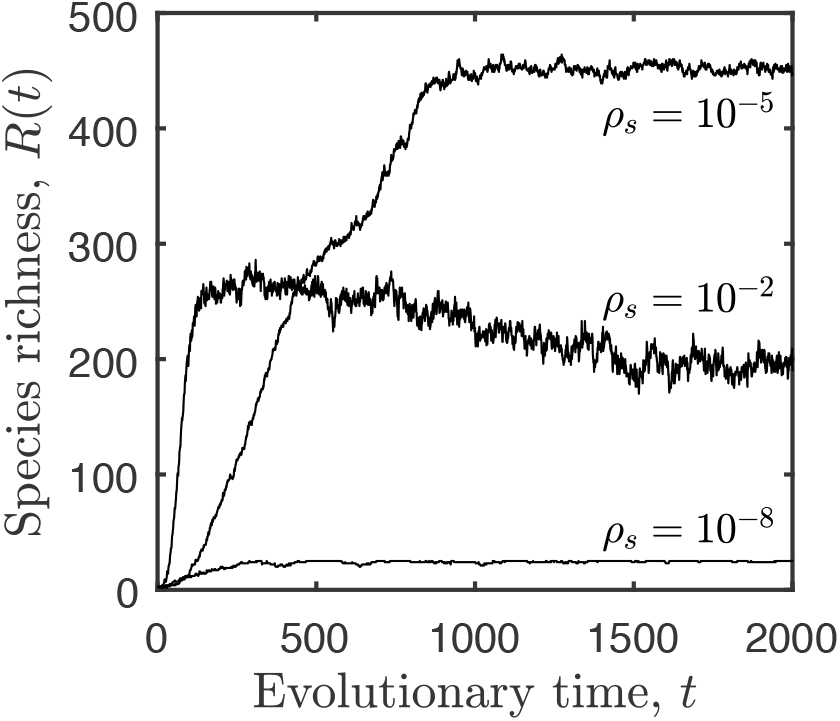
Example simulation dynamics for different dispersal rates, *ρ*_*s*_. When dispersal is low (*ρ*_*s*_ = 10^−8^), both the peak diversification rate, *σ*, and the quasi-equilibrium species richness, *R*\*, are low. When dispersal is intermediate (*ρ*_*s*_ = 10^−5^), the peak diversification rate is moderate and the quasi-equilibrium species richness is high. When dispersal is high (*ρ*_*s*_ = 10^−2^), the peak diversification rate is initially very high, but once all patches are occupied the species richness begins to fall due to competition among species that overlap in niche space, leading to lower quasi-equilibrium species richness. Parameters as in Fig. 4, with substitutable resources.

## Discussion

Our models confirm the intuitive prediction (Fig. 1) that maximal species richness should typically occur among organisms with intermediate dispersal (Fig. 3, 4). If dispersal rates are too low, then relatively little of the globally available area is found and occupied by the taxon, and the total number of unique species remains low. If dispersal rates are high, then this limitation on range spread ceases to be relevant, but now strong population mixing prevents divergence between occupied habitats; again, the total number of unique species remains low. It is therefore only at intermediate dispersal that most patches are occupied at a global scale, but the exchange of migrants remains low enough to permit divergence of resource use between patches. This pattern is qualitatively robust to a wide range of parameters, including variation in niche breadth, the number of available niches, carrying capacities, the number of patches, the relative speed of the ecological dynamics, baseline speciation rate, and time scales (Fig. S1–S8); it is therefore likely to be very general. Note that global species richness, being the outcome of speciation rates that operate over various areas, is not equivalent to speciation rate itself; if the same rate operates in a geographically restricted taxon, the outcome is fewer species than if the same rate applies over multiple geographic locations, thus the effect of dispersal definitely should be taken into account as it modifies the importance of regional versus global diversification (for a discussion of speciation in a biogeographic context see Schenk and Steppan 2018).

Notably, we find that global niche diversity can also peak at intermediate dispersal, but only when competition is for non-substitutable resources (e.g. food source and nest sites) as this leads to arbitrary non-overlapping associations between niche traits within each patch (Ashby et al. 2017). Under these conditions, intermediate dispersal produces different combinations of filled niches across patches, leading to high global niche diversity. Even though species sometimes disperse to other patches or new species arise locally, a priority effect (De Meester et al. 2016) tends to maintain established patterns within a patch and prevents the invasion of new species. Once the maximum number of species with non-overlapping traits has become locally established, any new or invading species will have to compete on multiple fronts, and will therefore be at a disadvantage compared to resident (non-overlapping) species. In contrast, competition for substitutable resources ultimately leads to high saturation of potential niches whenever a patch is occupied, even for very low dispersal. Although increasing dispersal leads to the colonization of new patches, it may not lead to a corresponding increase in niche diversity because the same set of niches can be filled in each patch.

Finally, we found that the peak diversification rate increased with dispersal (rather than peaking at intermediate dispersal). This is in part because high dispersal means new species will be more likely to disperse to a patch where they may be under weaker competition than with their ancestor, but also because high dispersal leads to rapid colonization of all patches and hence a high number of locally-available niches for species to fill. Indeed, when a patch is newly colonized, there is likely to be a short period of rapid diversification (i.e. an adaptive radiation) as species partition the available niches, followed by a reduction in the diversification rate (Crouch and Ricklefs 2019).

As a whole, our model has potential to unify previous findings, as it shows why it is possible to find both increasing (Owens et al. 1999) and decreasing (Jablonski 1986) effects of dispersal rate on speciation. These findings may reflect distinct but often co-occurring processes, visually depicted as taxa potentially residing on different slopes of a unimodal (humped) shape. On the left slope (Fig. 1), more dispersal leads to higher regional species richness, as speciation is limited by the rate at which lineages spread to new areas in which speciation can occur. On the right slope, dispersal has surpassed the point where it leads to maximum speciation; further increases in dispersal lead to dynamics where frequent exchange of migrants maintains few but very widely distributed species. Obviously, it remains a challenge to make a precise statement of the location of a taxon’s dispersal ability with respect to one that would maximize diversification, but it appears plausible that taxa with a currently globally restricted range might have failed to disperse to establish where they in principle could – or they disperse too infrequently to overcome the fact that other taxa have taken the niche and prevent new invasions via a priority effect (De Meester et al. 2016), while very widespread species ranges (e.g. those of high-latitude Sylvia warblers, (Böhning-Gaese et al. 2006)) could indicate the opposite and permit too widespread gene flow for speciation to occur readily.

What do our results add to Haldane’s apocryphal quip (Farrell 1998)? They definitely allow speculating on a potential causality: beetles have wings but they use them relatively infrequently, and over shorter distances, compared with many other volant organisms. Obviously, it is difficult to go from a natural history fact to a quantitative statement of beetles being located, of all taxa, closest to the peak of Fig. 1, especially since the relevant variation in the number of niches available to be filled may be taxon-specific. Indeed, Hymenoptera may challenge the status of Coleoptera as the most speciose order as a result of the prevalence of the parasitic lifestyle in hymenopterans, leading to specialism in host use (Forbes et al. 2018). Nor do our results claim dispersal to be the only causal factor for Coleoptera: suggested drivers of species richness in this taxon include feeding on angiosperms (Farrell 1998) to variation in the ability to resist extinction (Smith and Marcot 2015); moreover, not all beetles are reluctant fliers (Brown et al. in press). We would rather argue that to achieve the status of exceptional diversity, all relevant factors need to come together in a manner that jointly maximizes the diversification rate. Our model confirms earlier verbal arguments (Kneitel and Miller 2003, Price and Wagner 2004, Claramunt et al. 2012, Agnarsson et al. 2014) that suitable dispersal rates form one such factor, and we would welcome an empirical effort to classify species according to their chances to complete long-distance dispersal attempts to new sites.

In this quest, the relevant aspect of dispersal cannot be boiled down to a simple dichotomy between wings (with a suitable rate of using them) vs. winglessness; any type of suitably infrequent colonization of new areas is of interest. The hyperdiverse genus of flightless weevils *Trigonopterus*, inhabiting the Sunda Arc in southeast Asia, provides a thought-provoking example. Reconstructions of the colonization history of the various islands in this archipelago show repeated arrivals of different *Trigonopterus* clades, with e.g. Sulawesi receiving new extant clades at intervals of several million years (Tänzler et al. 2016). Especially since this genus occasionally manages to cross Wallace’s line, an aquatic mode of dispersal (rafting) is suspected for this genus’ long-distance colonization events (Tänzler et al. 2016). Direct experimental evidence is available for *Pachyrhynchus* (also a flightless weevil), where eggs and larvae survive oceanic conditions inside floating fruit in both laboratory and field conditions (Yeh et al. 2018).

While we appreciate the challenges of quantifying the frequency of dispersal attempts, especially when many colonization attempts fail due to stochasticity or priority effects, our modelling can also highlight further avenues of work. Both theoretically and empirically, a clear next step would be to add a consideration of any temporal trends (over evolutionary timescales) in dispersal ability. Beetle speciation appears to be particularly sped up in cases where lineages that first dispersed to new areas subsequently lost the ability to fly (Ikeda et al. 2012) (for evidence that colonization ability of beetles covaries with flight ability see (Iversen et al. 2017)). Patchy environments (e.g. archipelagos for terrestrial organisms) provide particularly intriguing food for thought in this context, since the initial colonization requires dispersal, but this may be followed by selection to reduce dispersal rates (or abilities) if large amounts of matrix habitat between suitable sites make dispersal risky at the individual level (Diamond 1981, Shaw et al. 2014). Simplified island ecosystems that lack predators may concurrently lead to loss of flight in organisms that would use flight to flee. In birds, not only are flightless species commonly restricted to oceanic islands (Feduccia 1980, McNab 1994); flying birds residing on such islands also show a repeatable pattern of reduced investment in flight-related morphology (Wright et al. 2016). Global diversification of a taxon will be particularly enhanced if post-colonization reductions in dispersal ability (Wright et al. 2016) promote speciation (Fulton et al. 2012, Ikeda et al. 2012). However, since extinction rates may simultaneously increase (Shaw et al. 2014), the net effect remains to be investigated.

Like any model of evolutionary ecology, ours makes a compromise between generality (with its messages hopefully applying across many taxa) and the precision and inclusiveness regarding many phenomena that might vary in their taxon-specific importance. By leaving much detail (regarding e.g. biogeography or genetic architecture) outside the model, we have kept the focus on how resource competition can lead to contrasting distributions of niches as a function of dispersal, and in turn the likely consequences for species richness and diversification rates. Even within our chosen focus, knowledge gaps remain. Our model assumed there were only two niche dimensions, but our findings should extrapolate to multiple niche dimensions due to the overlapping/non-overlapping patterns that these different resource competitions are known to produce (Ashby et al. 2017). In reality, niche space will consist of both substitutable and non-substitutable resources, which means that species will densely-pack regions of the niche space representing substitutable axes and sparsely-pack those representing non-substitutable axes. While we expect no qualitative impact on the patterns for species richness as both resource types cause a peak at intermediate dispersal, adding substitutable resource axes will lead to a greater proportion of filled niches at both the local and global levels, whereas the addition of non-substitutable should have the converse effect.

The motivation for our work was the curious lack of theoretical models testing the prediction of a hump-shaped relationship between dispersal and species richness, and as a result our study has implications for both empiricists and theoreticians. For empiricists, we have shown this relationship to be robust to a wide range of modelling assumptions, which suggests the fundamental concept is likely to be fairly general. But we have also made the prediction that the qualitative relationship between dispersal and niche diversity in the metacommunity depends on the mode of resource competition, which should be tested alongside the dispersal-species richness prediction. For theoreticians, future, more complex models of the dispersal and speciation processes should add nuance to the relationship we found, hopefully exploring whether the verbal model of dispersal and speciation remains as mathematically sound as our work, as a first step in this direction, shows them to be. Our work therefore sets the stage for further theoretical work on more complex speciation processes and temporal (evolutionary) changes in dispersal ability, taking into account the potentially contrasting (Mouquet and Loreau 2003) effects of dispersal on local, regional and global diversity patterns.

## Data accessibility

Simulation code is available in the *Supplementary material*.

## Acknowledgements

BA is supported by the Natural Environment Research Council (grant no. NE/N014979/1). AKS is supported in part by the National Science Foundation. HK acknowledges the support of the Swiss National Foundation (grant no. 31003A_163374).

## Author contributions

A.K.S. and H.K. conceived the study and B.A. carried out the modelling work. All authors contributed equally to writing the manuscript. The authors declare no conflicts of interest.

---

**Figure S1.**
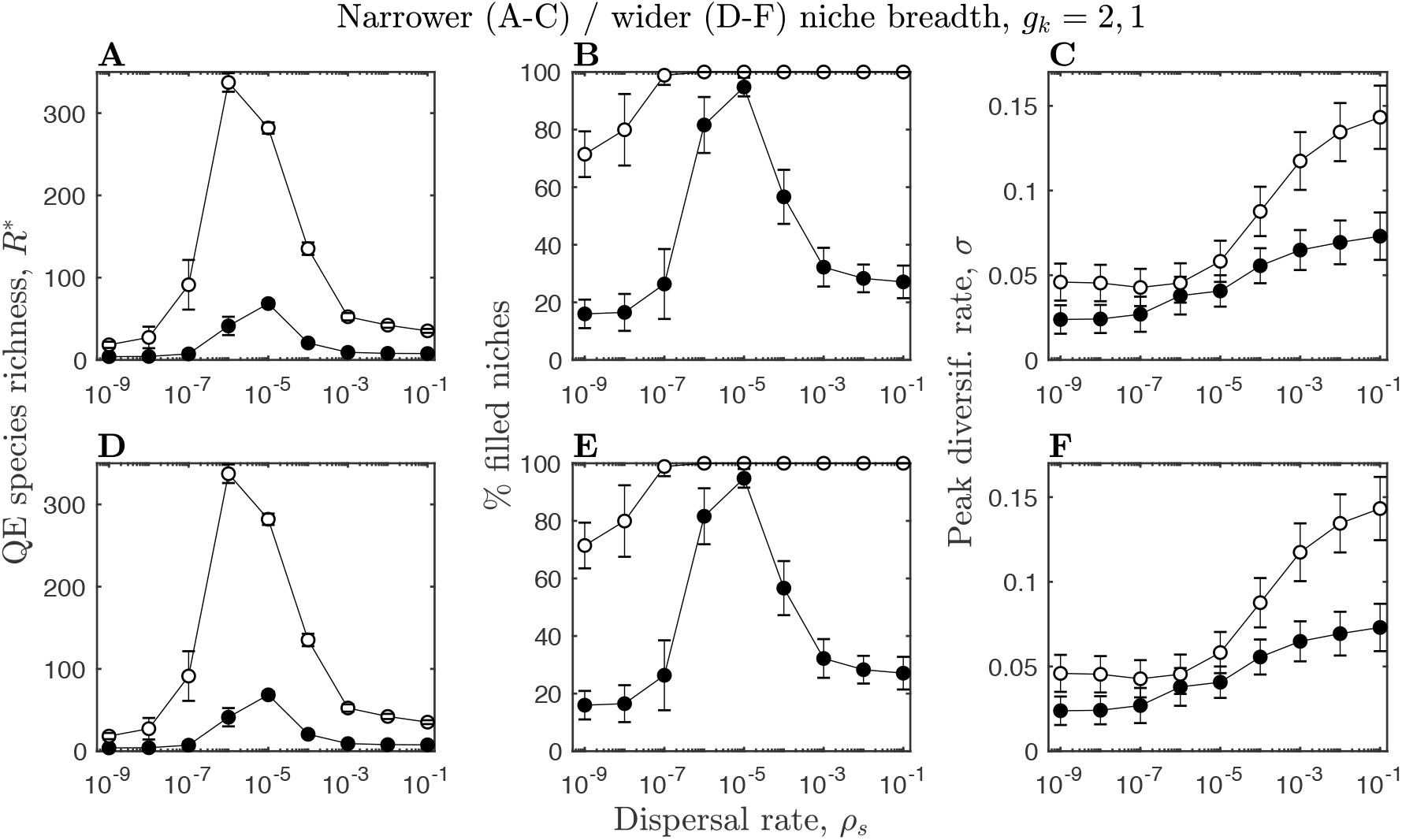
Parameter sensitivity analysis for narrower (top row, *g*_*k*_ = 2) and wider (bottom row, *g*_*k*_ = 1) niche breadths. Figure details and remaining parameters as described in Fig. 4.

**Figure S2.**
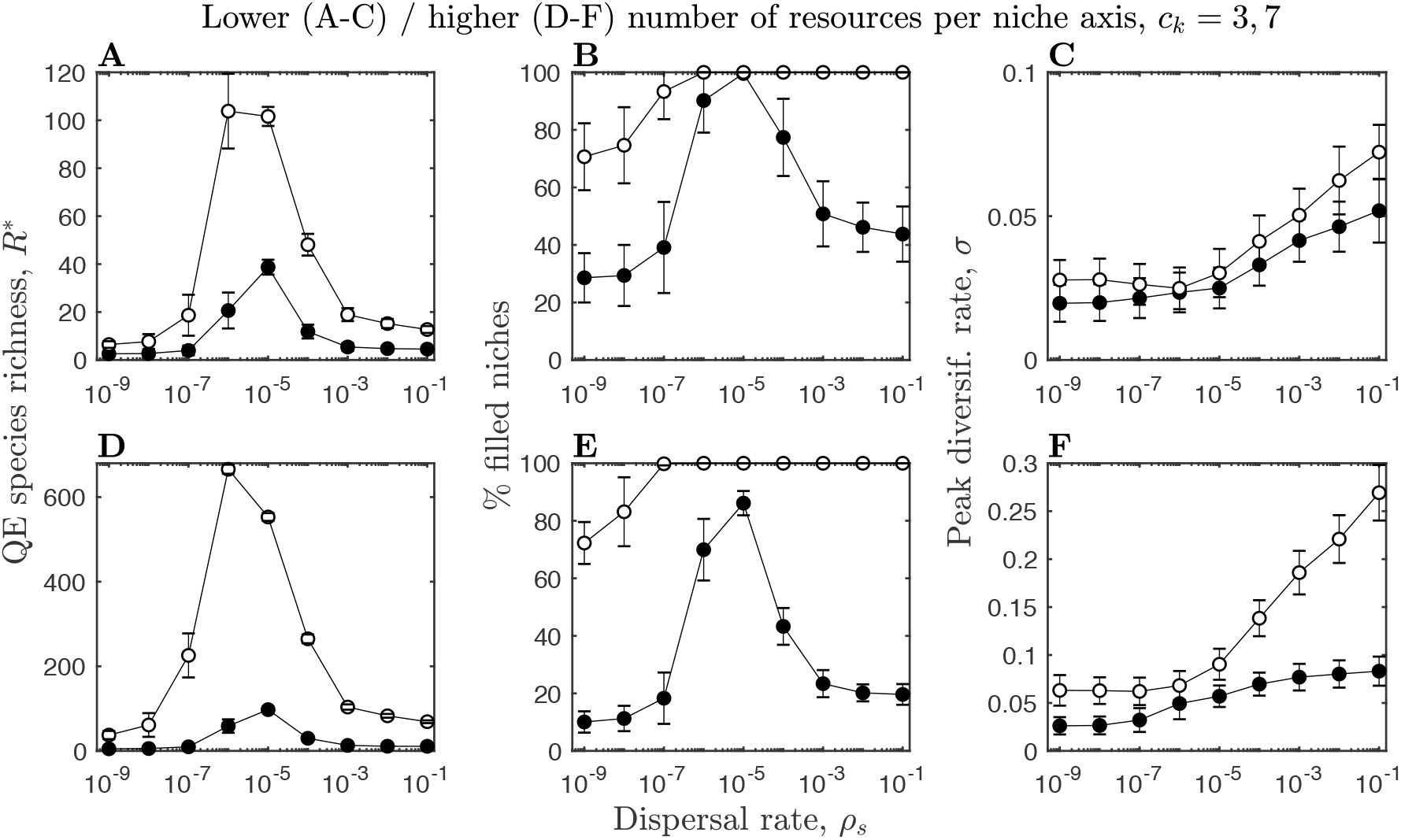
Parameter sensitivity analysis for fewer (top row, *c*_*k*_ = 3) and more (bottom row, *c*_*k*_ = 7) resources per niche axis. Figure details and remaining parameters as described in Fig. 4.

**Figure S3.**
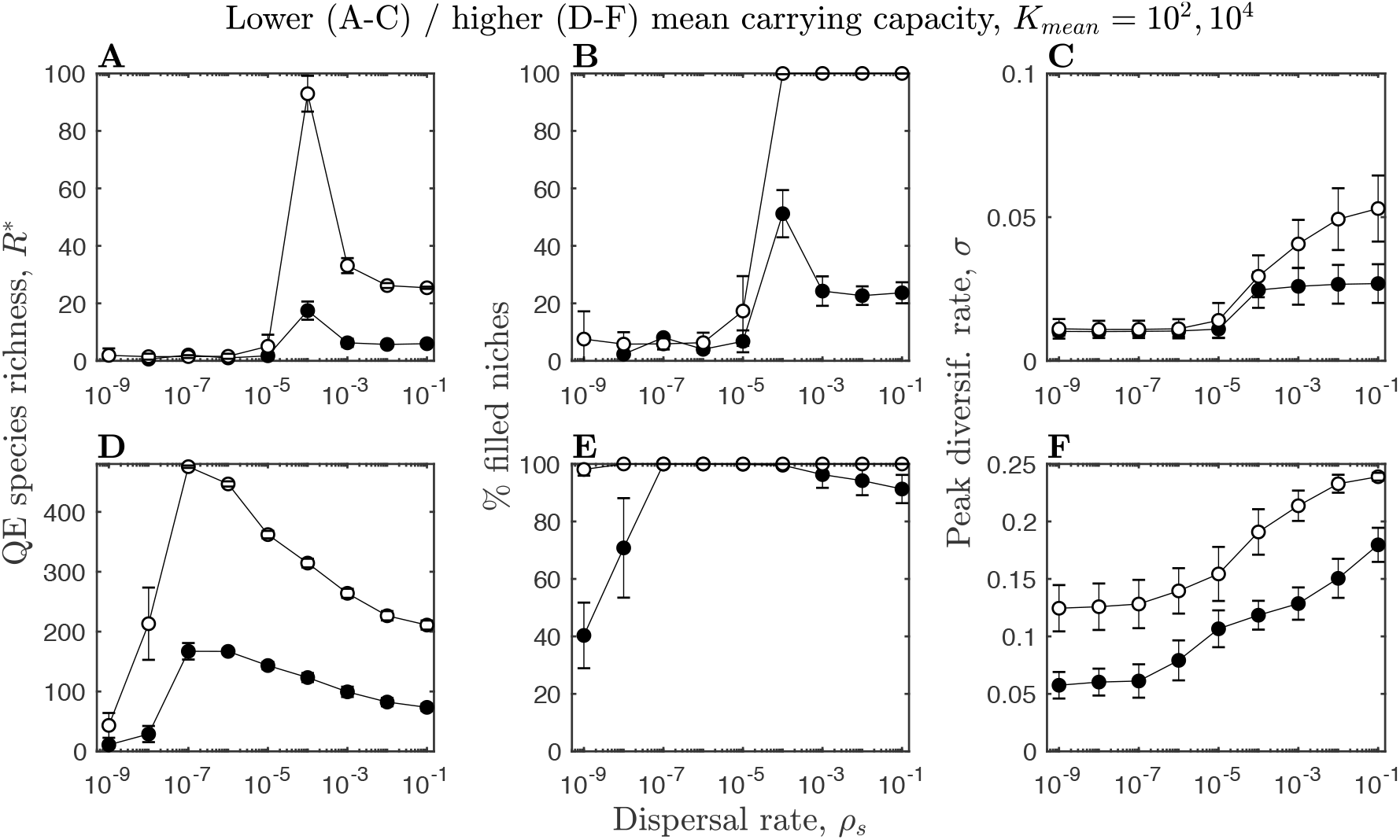
Parameter sensitivity analysis for lower (top row, *K*_*mean*_ = 10^2^) and higher (bottom row, *K*_*mean*_ = 10^4^) mean carrying capacities. Figure details and remaining parameters as described in Fig. 4.

**Figure S4.**
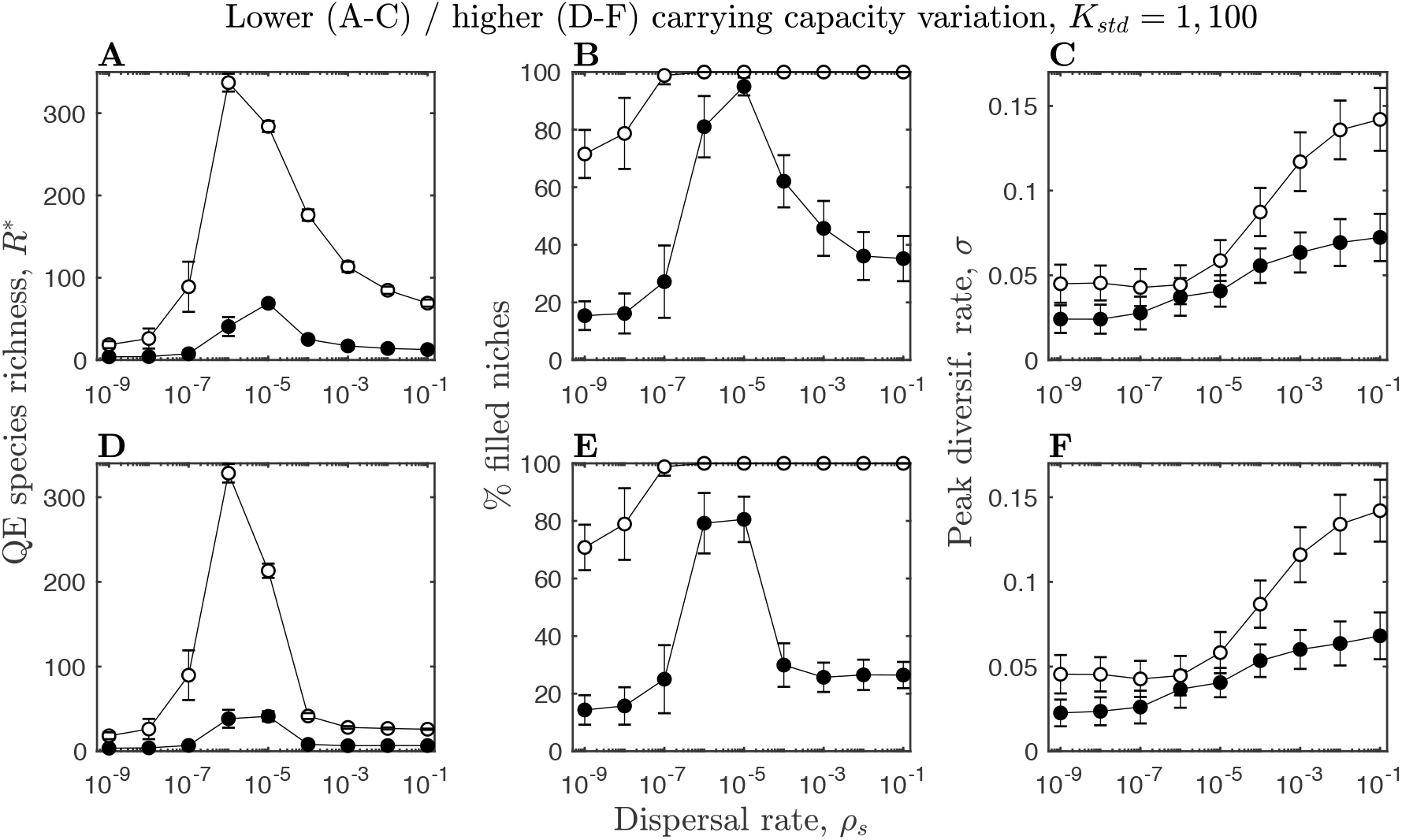
Parameter sensitivity analysis for lower (top row, *K*_*std*_ = 1) and higher (bottom row, *K*_*std*_ = 100) standard deviations in carrying capacities. Figure details and remaining parameters as described in Fig. 4.

**Figure S5.**
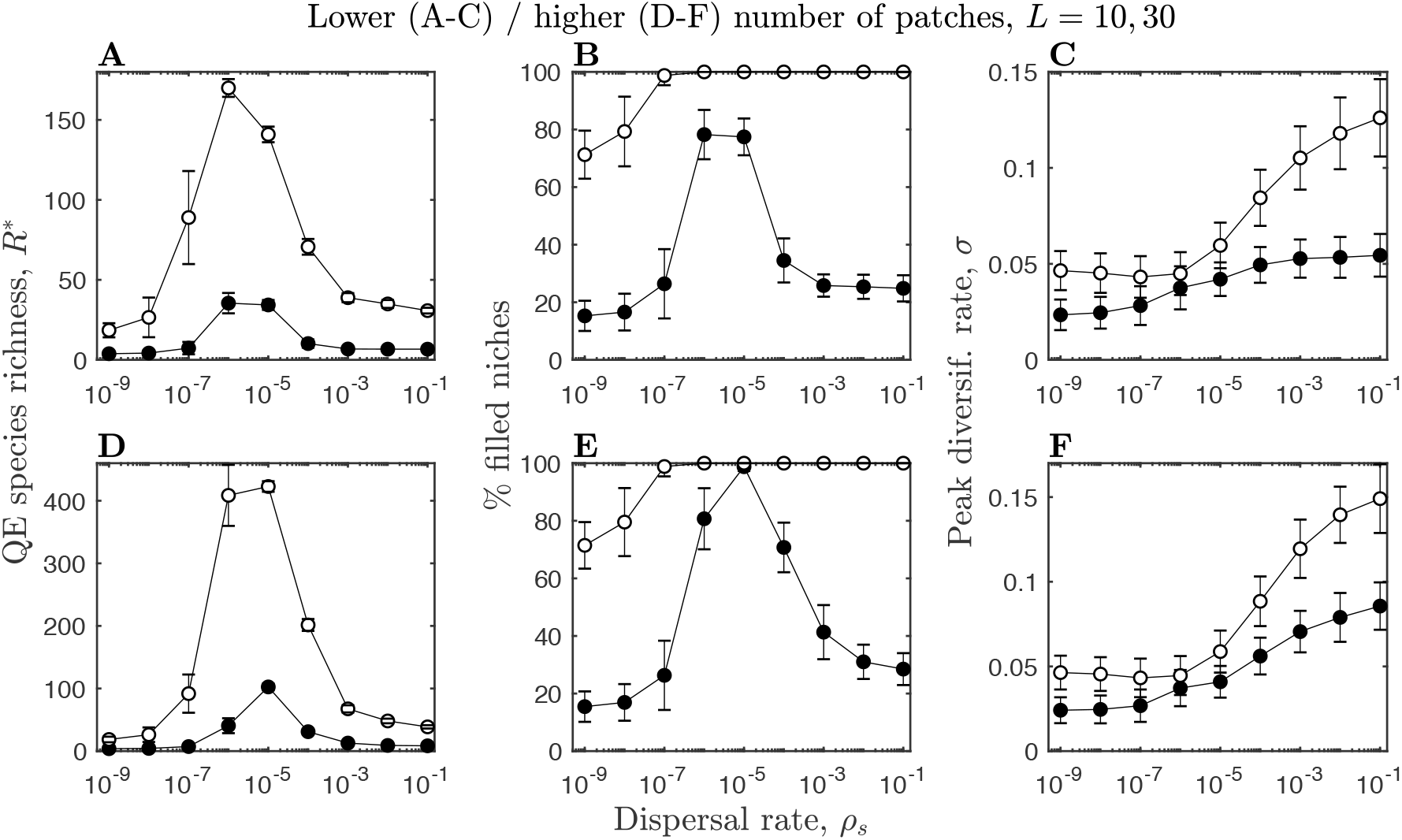
Parameter sensitivity analysis for fewer (top row, *L* = 10) and more (bottom row, *L* = 30) patches. Figure details and remaining parameters as described in Fig. 4.

**Figure S6.**
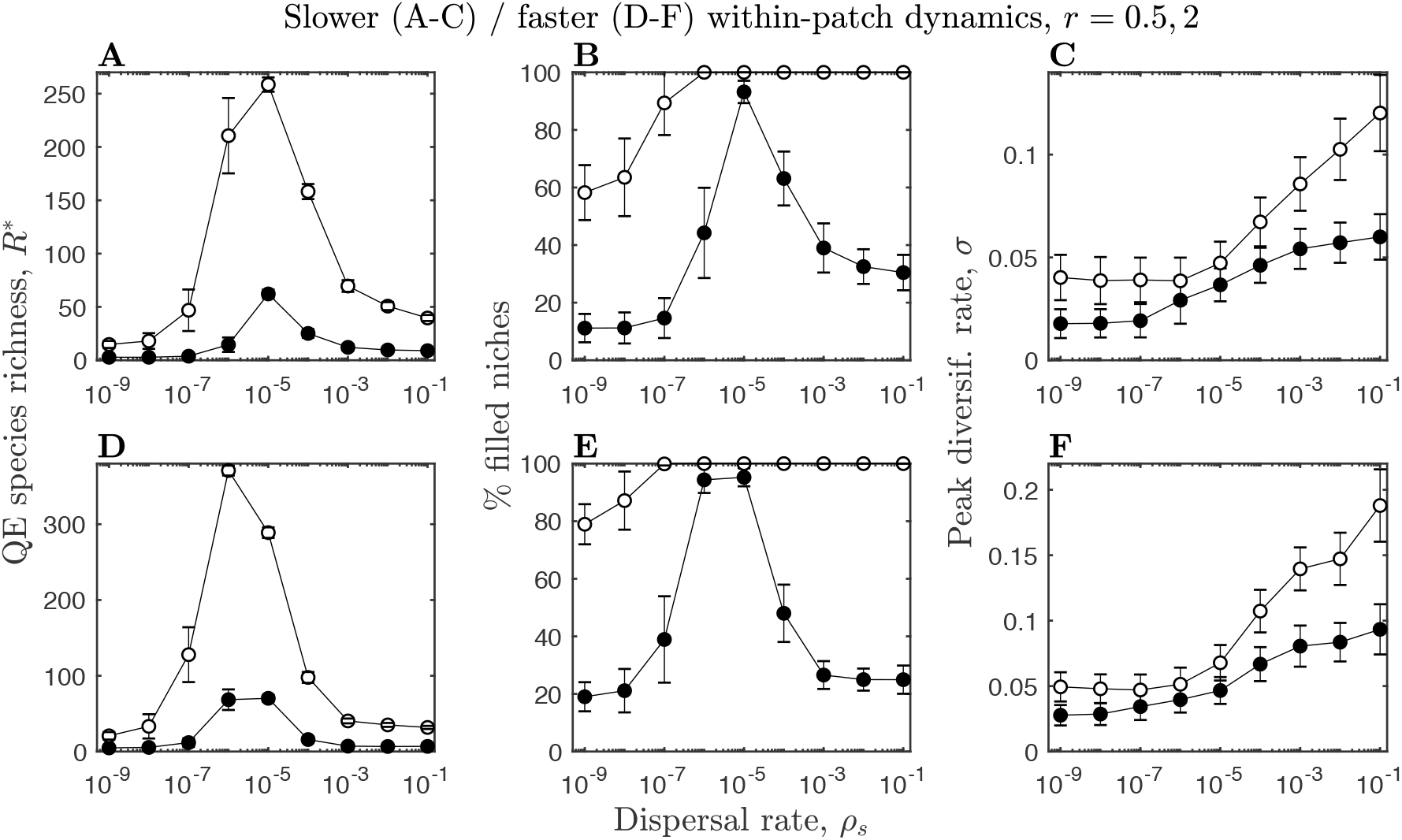
Parameter sensitivity analysis for slower (top row, *r* = 0.5) and faster (bottom row, *r* = 2) within-patch dynamics. Figure details and remaining parameters as described in Fig. 4.

**Figure S7.**
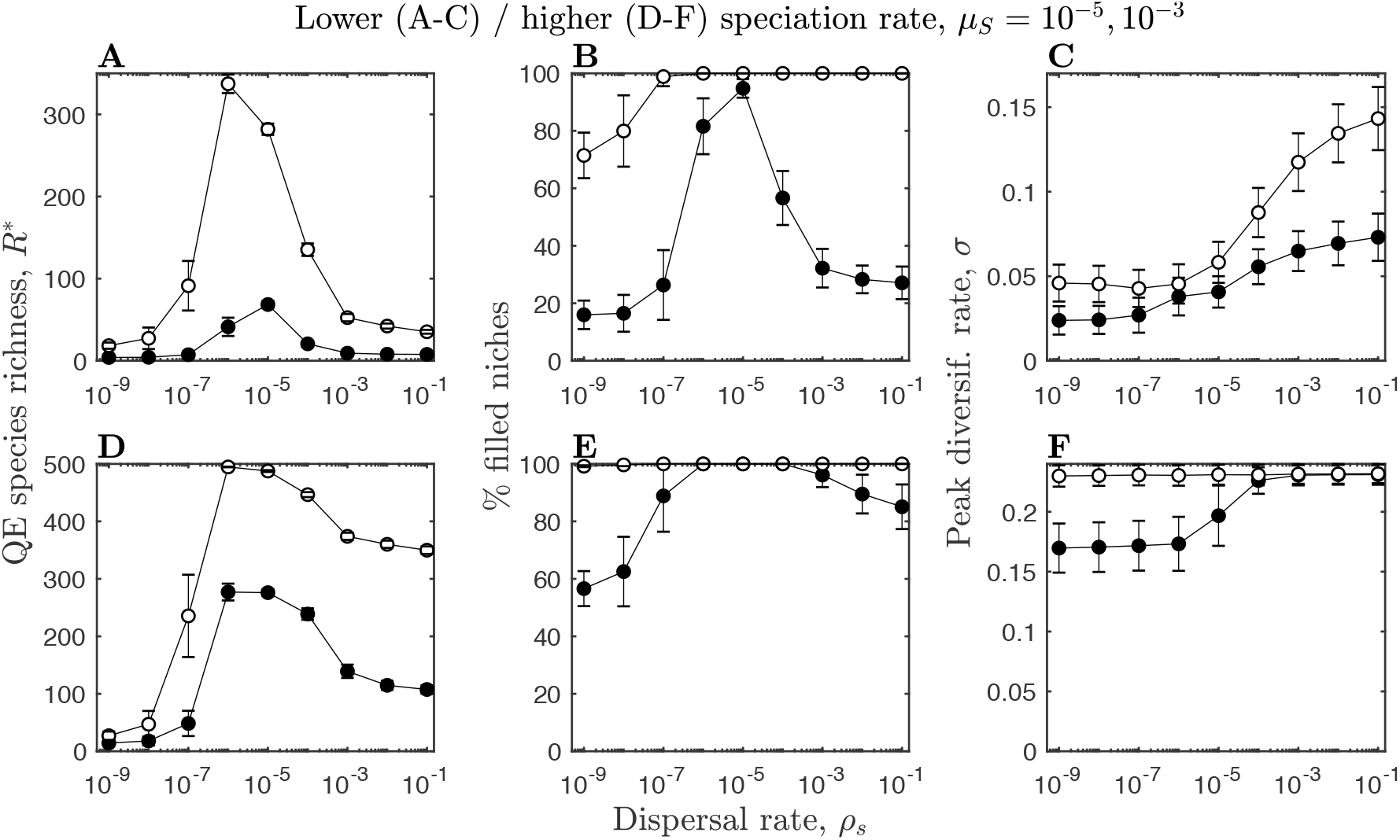
Parameter sensitivity analysis for slower (top row, *μ*_*S*_ = 2 × 10^−4^) and faster (bottom row, *μ*_*s*_ = 2 × 10^−3^) speciation rates. Figure details and remaining parameters as described in Fig. 4.

**Figure S8.**
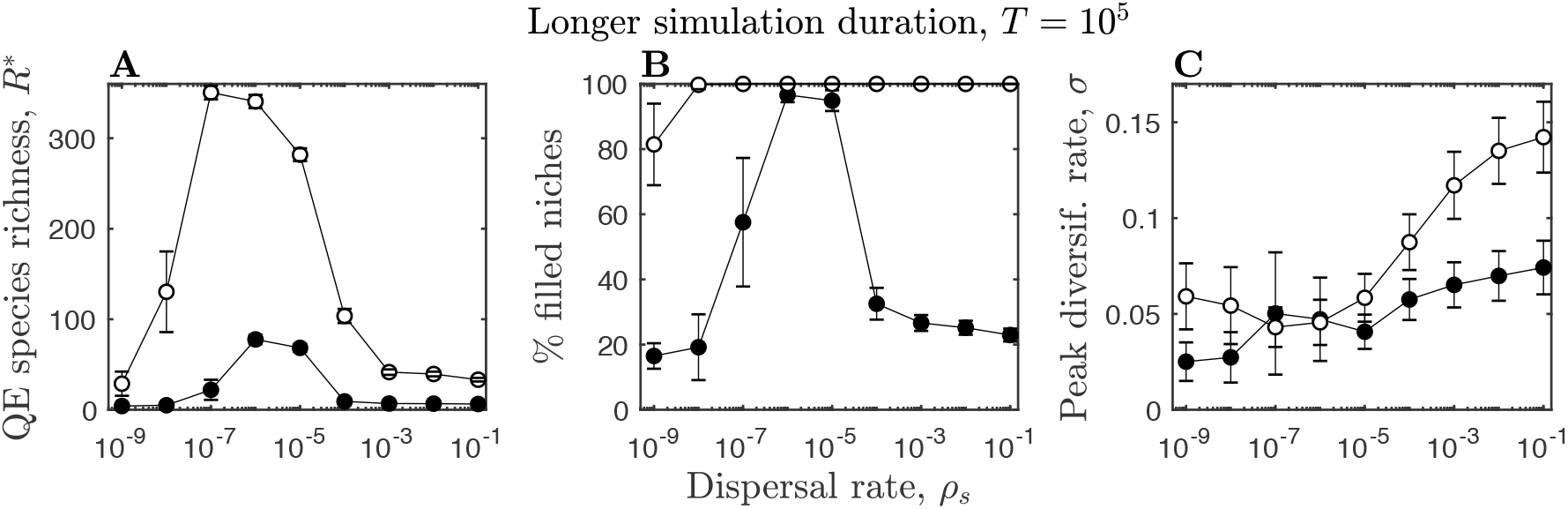
Parameter sensitivity analysis for longer simulation durations (*T* = 10^5^). Figure details and remaining parameters as described in Fig. 4.

## Source code

~~~
function [species_total,snapshots] =
dispersal_speciation(max_time,r,Kmean,Kstd,rho,mu,epsilon,gk,ck,L,niche_typ
e,sample_points)
~~~

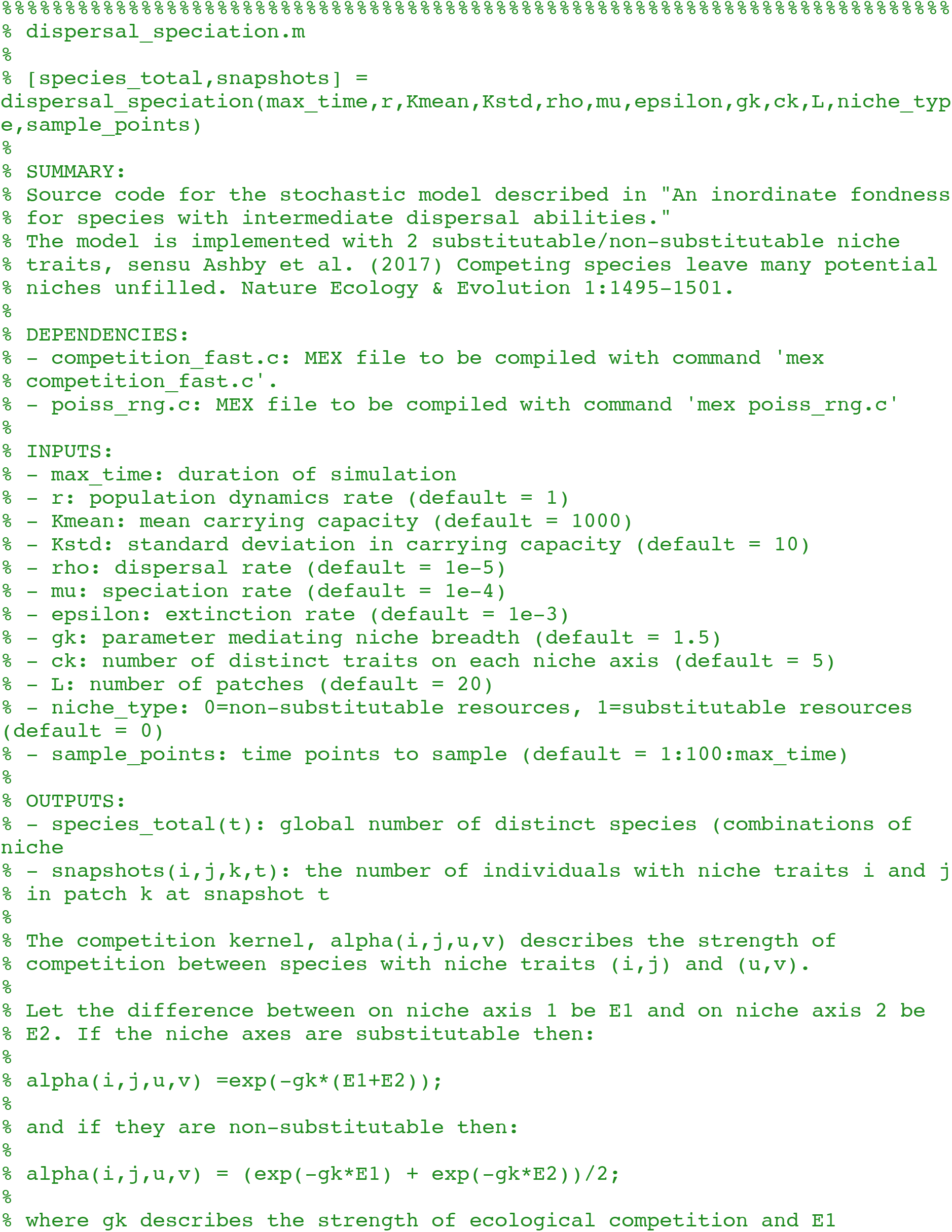
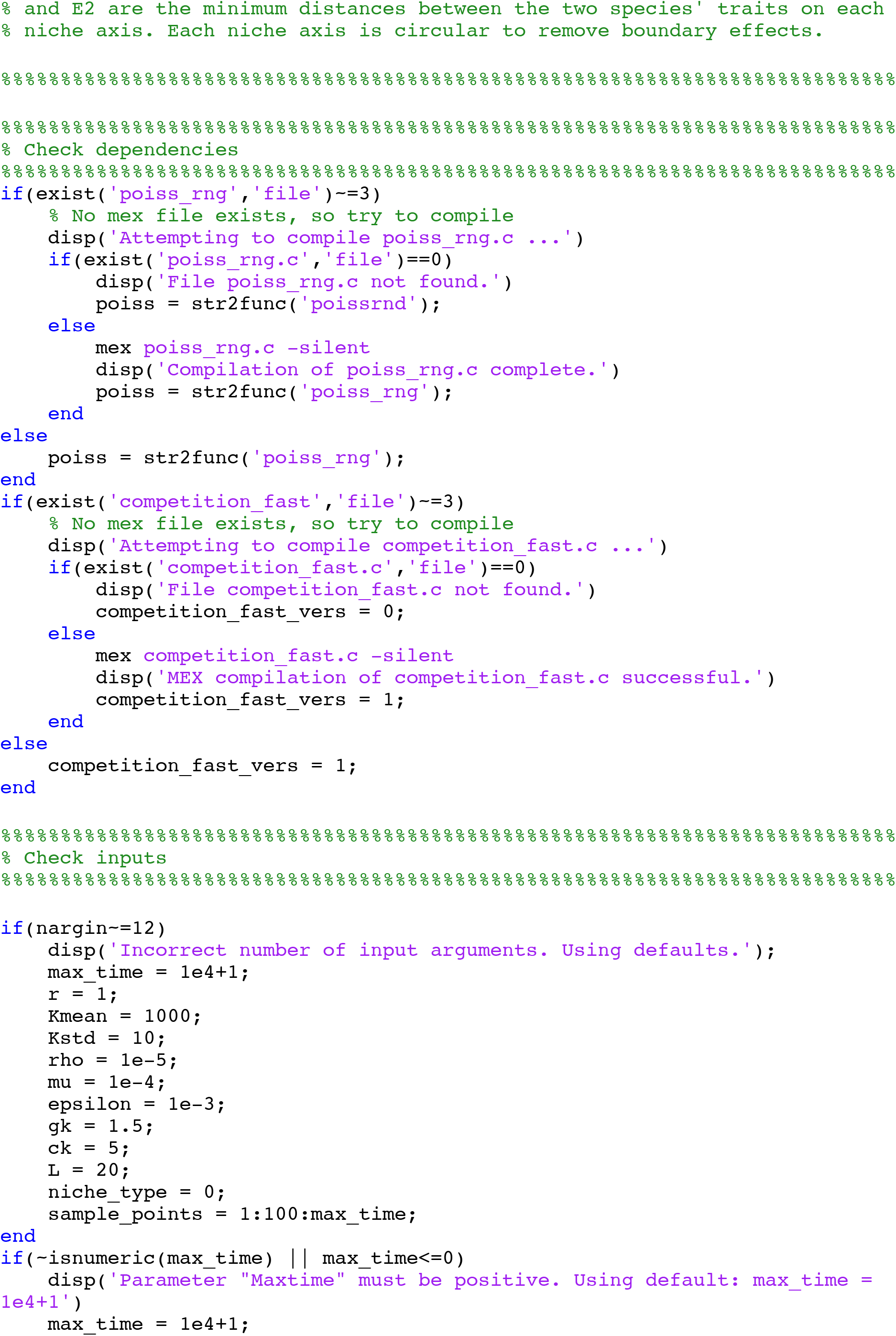
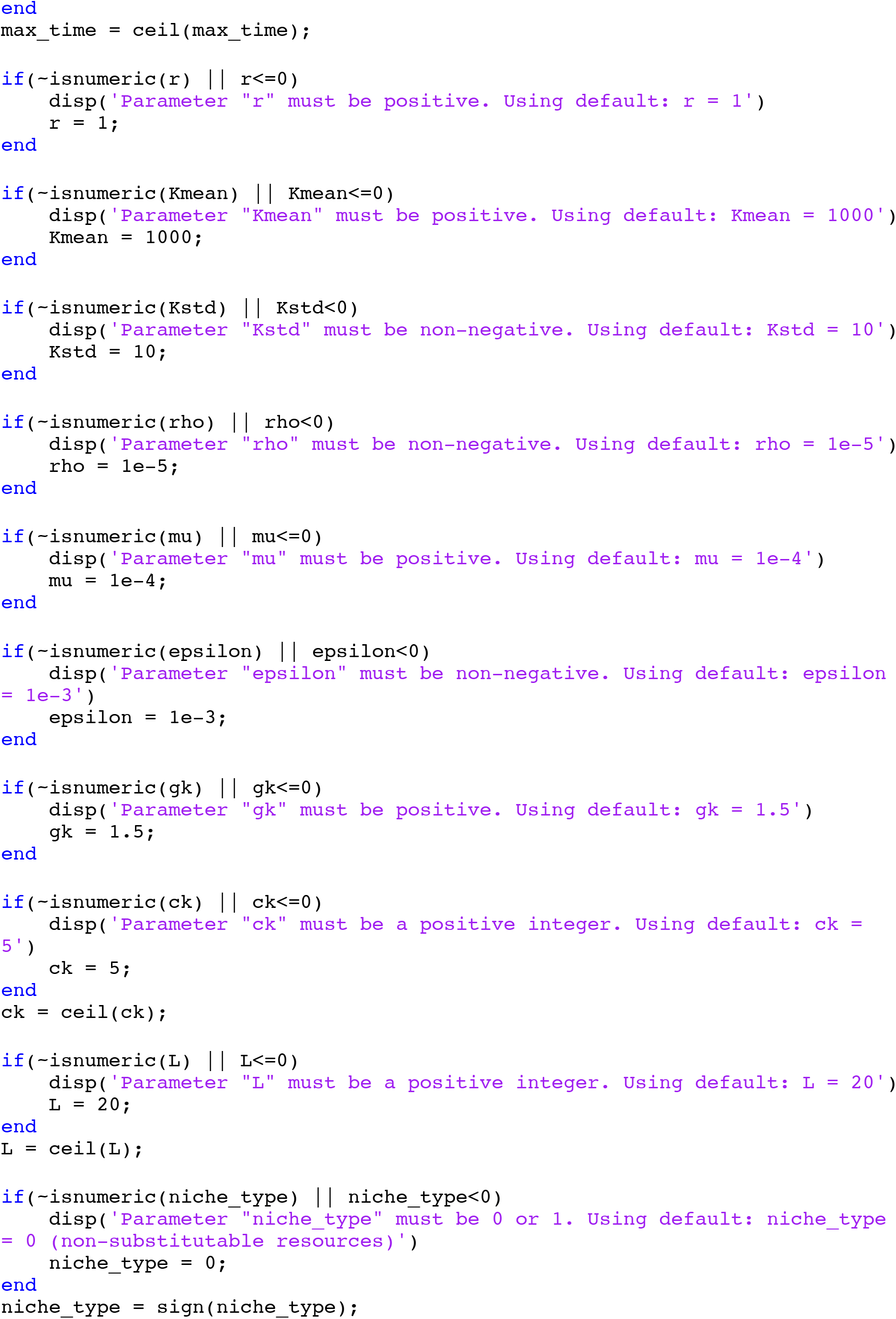
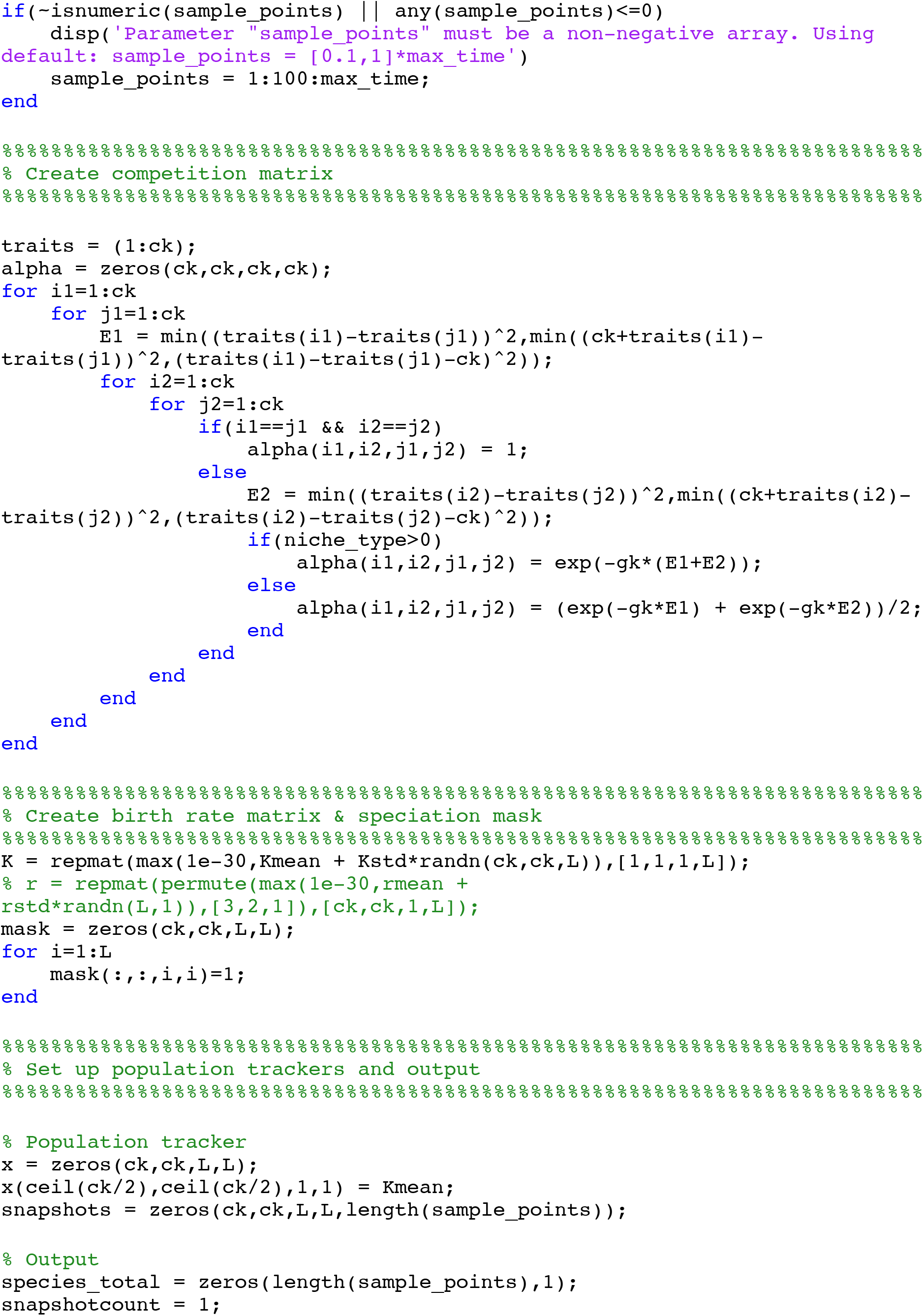
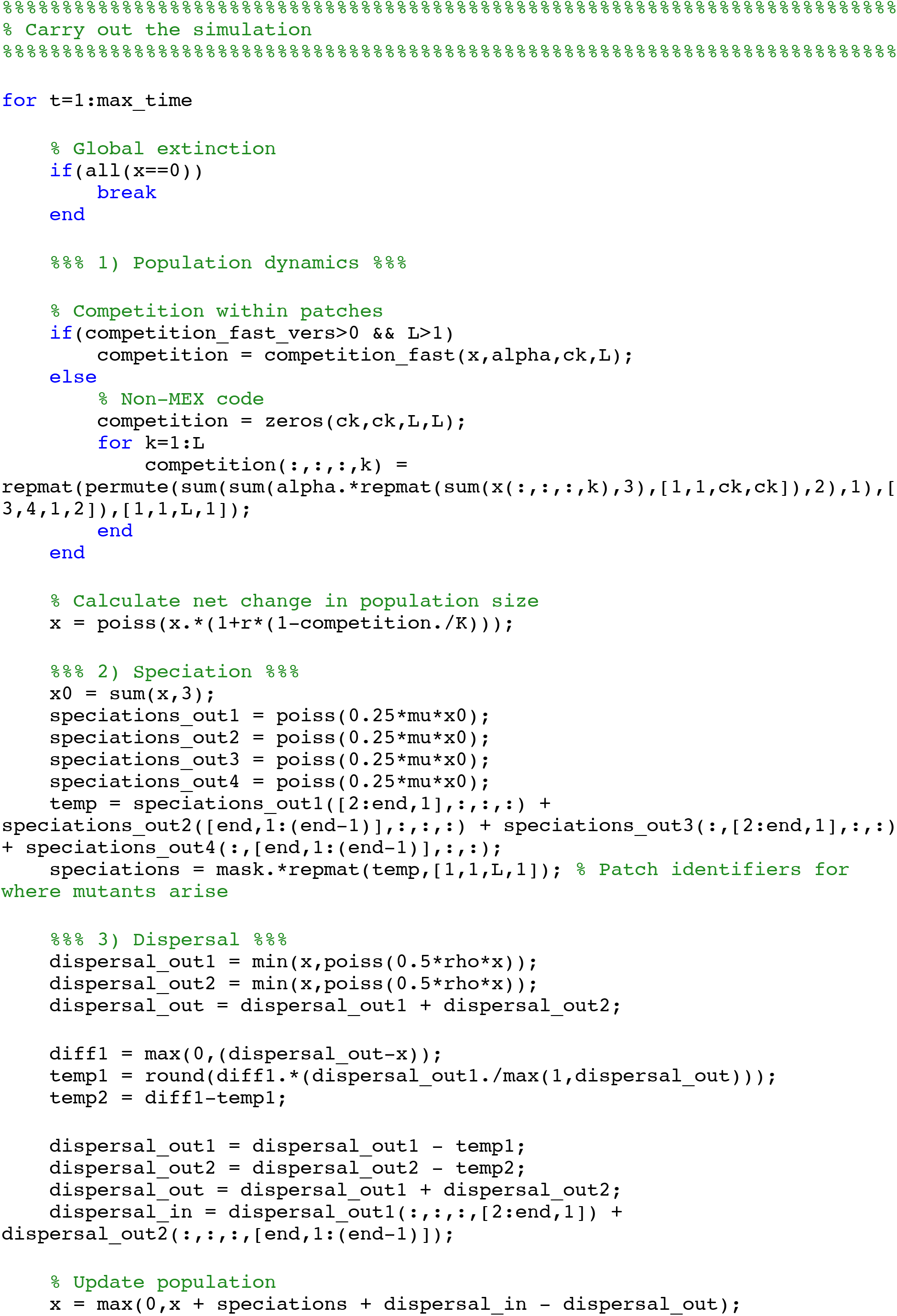
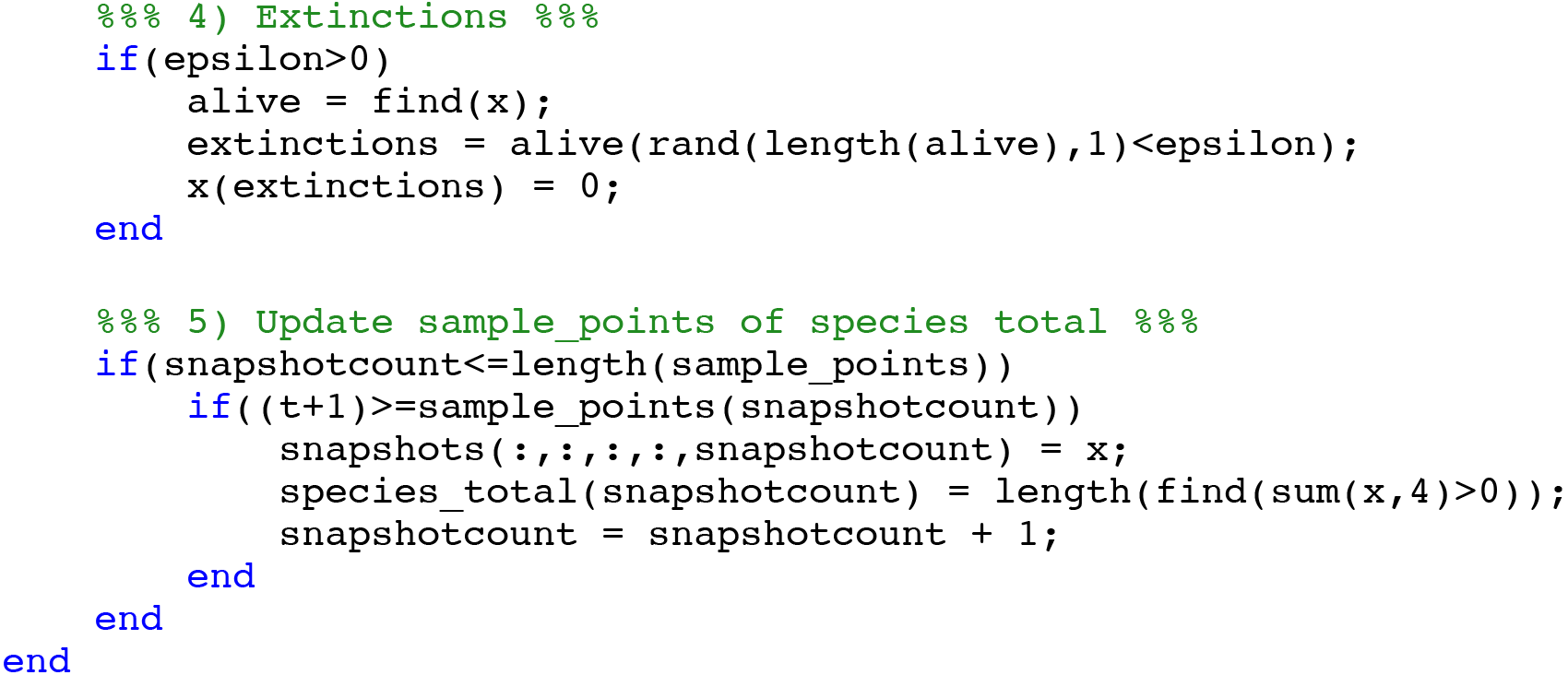

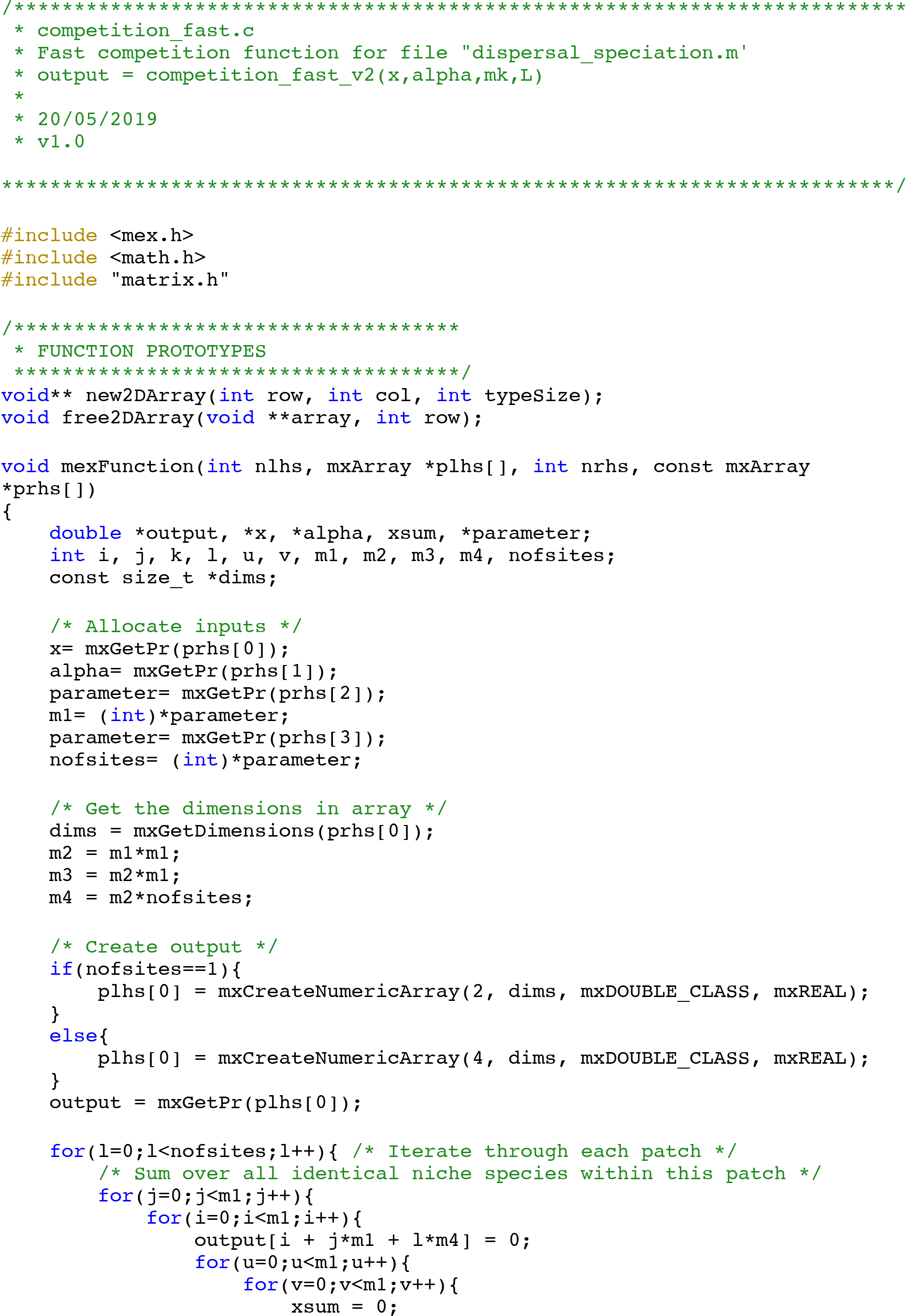
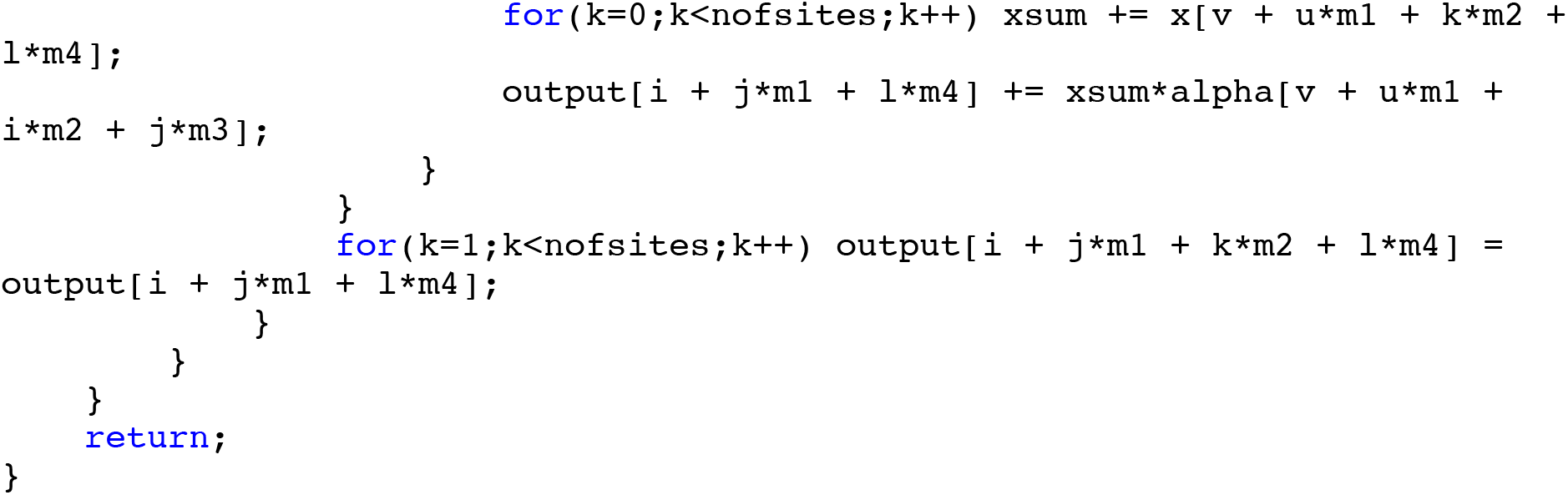

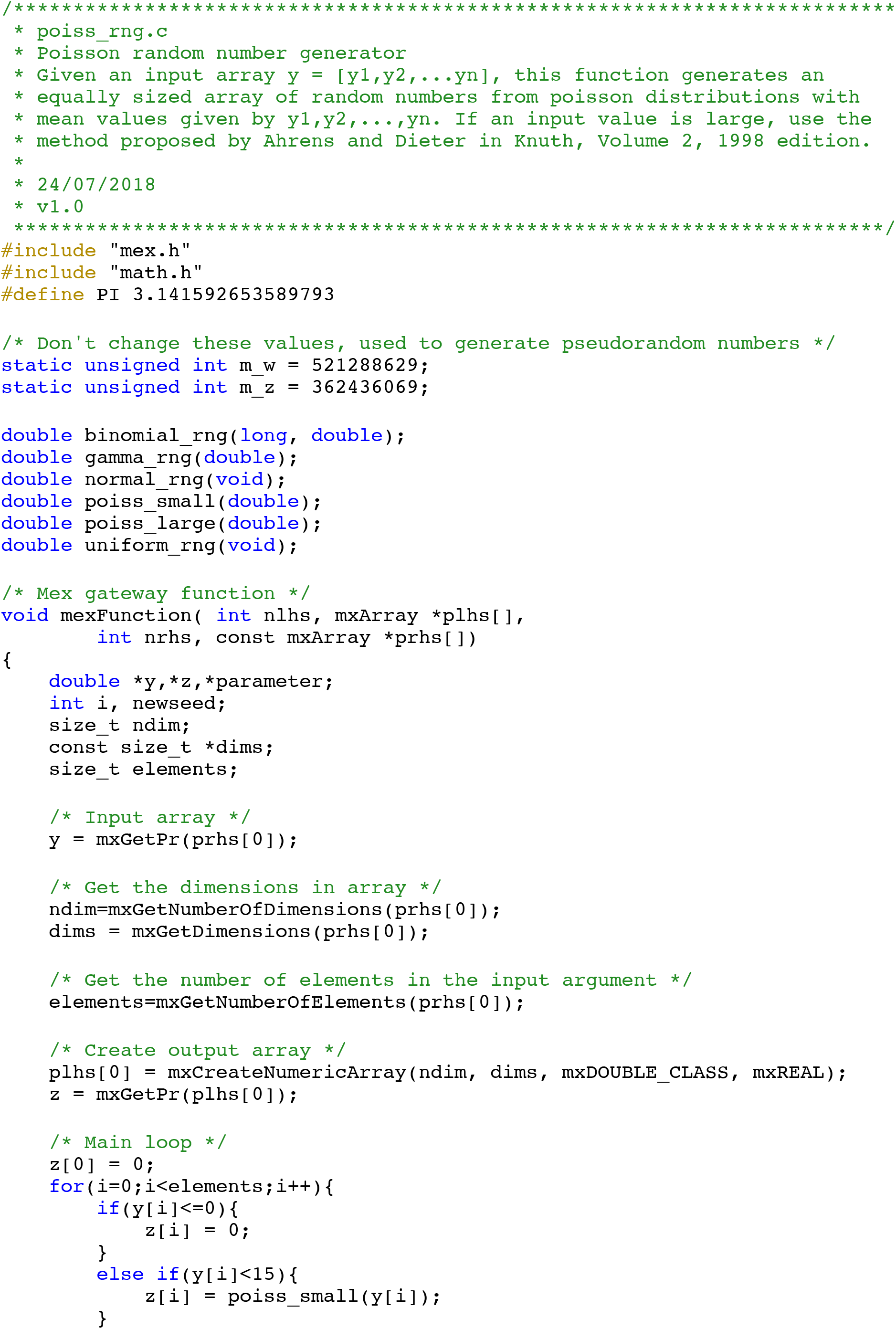
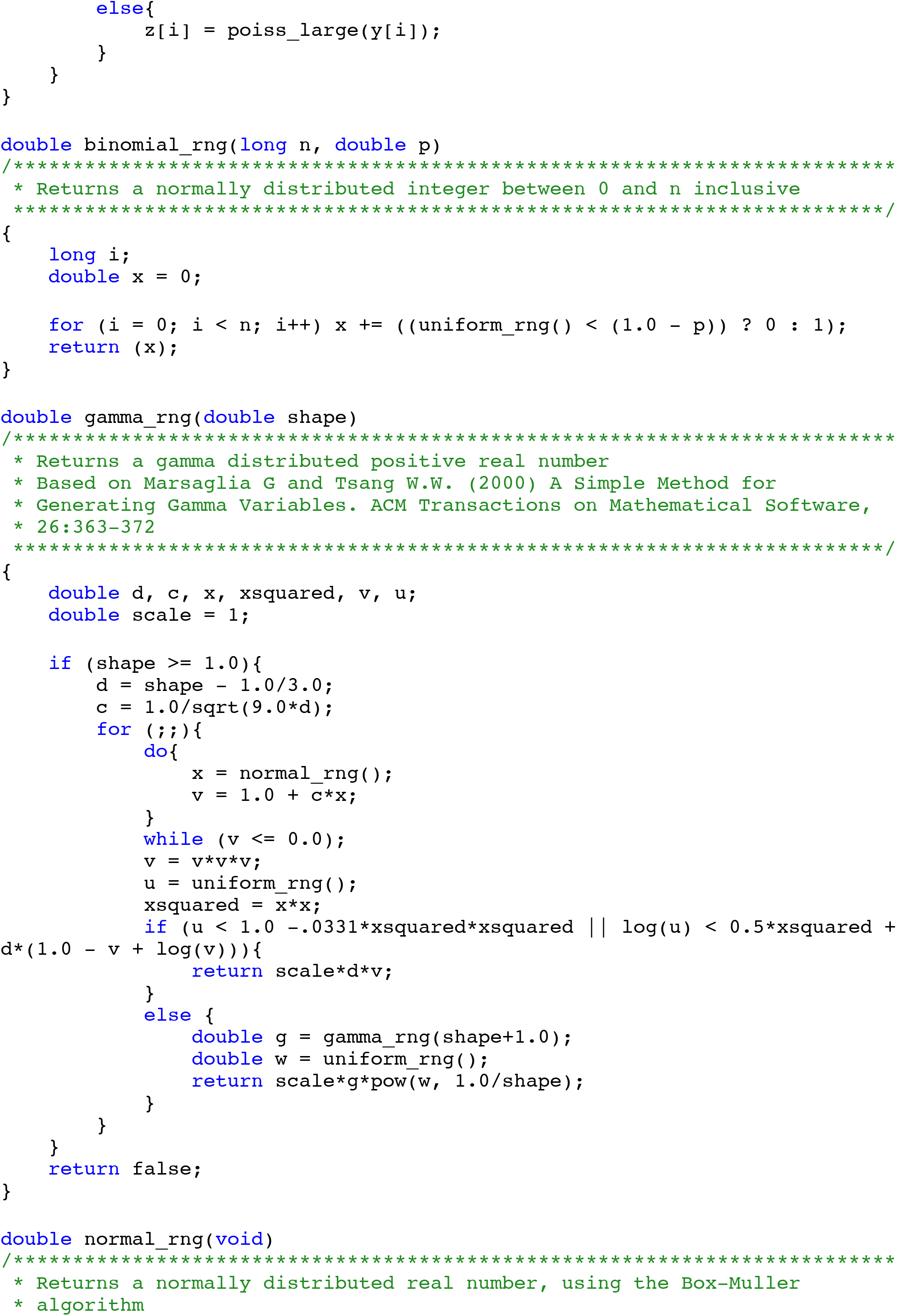
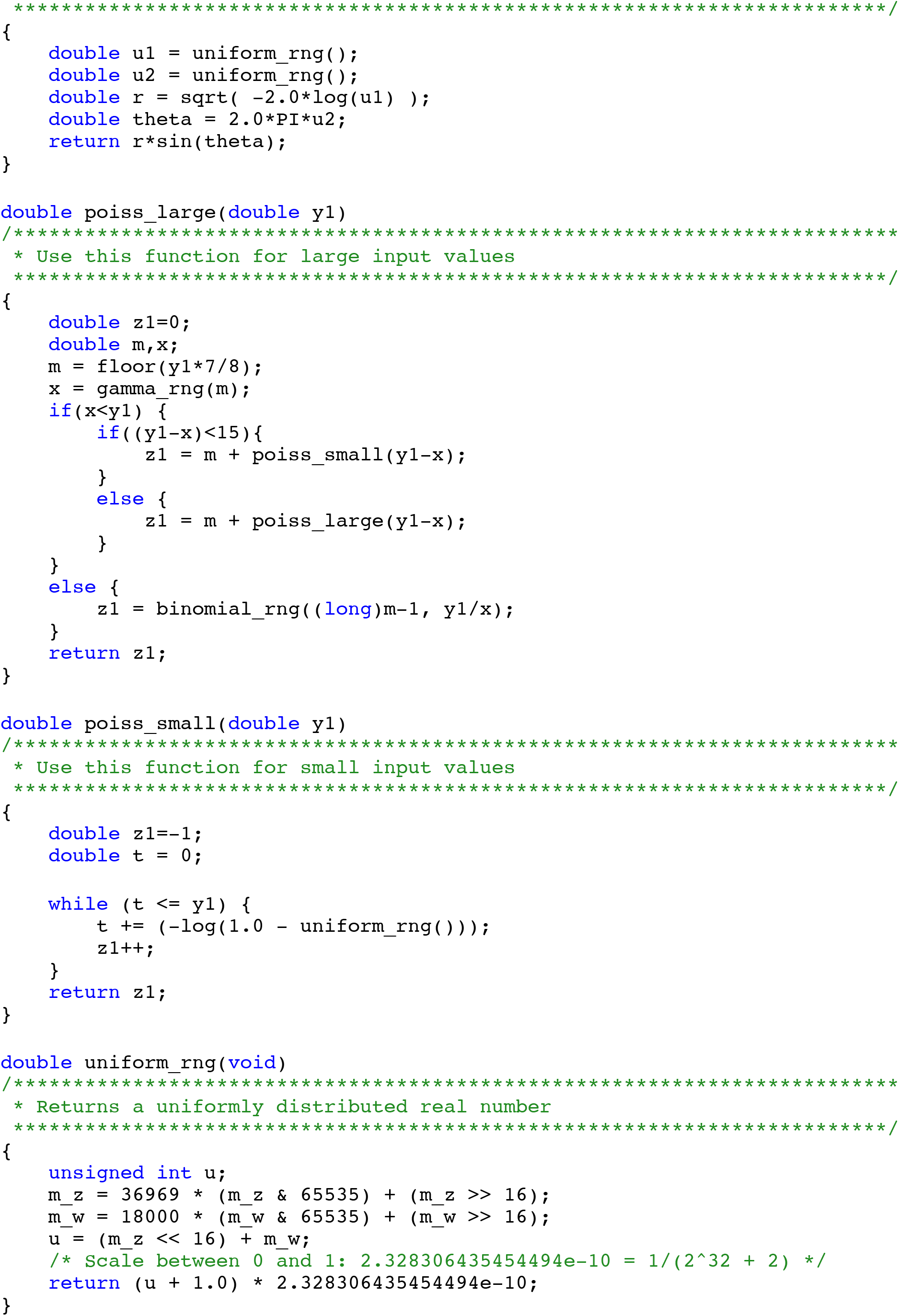

