## Supplementary material for "An Inordinate Fondness for Species with Intermediate Dispersal Abilities": Fig. S1-S8

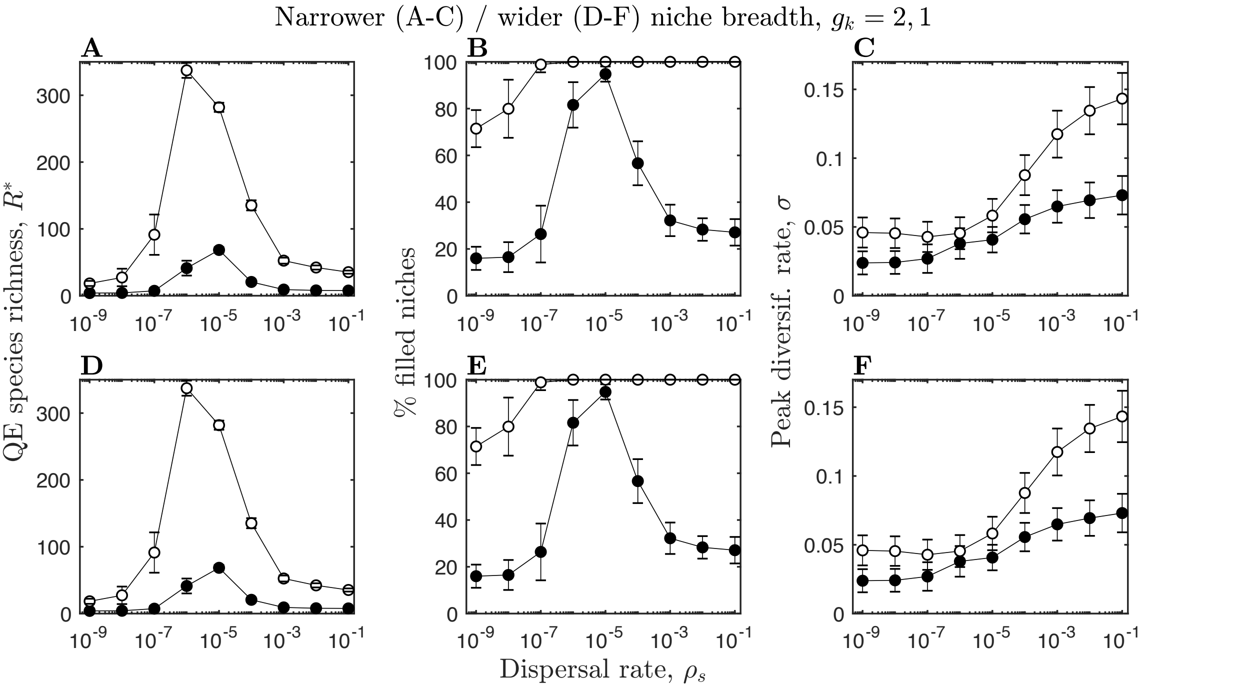


Figure S1 – Parameter sensitivity analysis for narrower (top row, $g_k = 2$) and wider (bottom row, $g_k = 1$) niche breadths. Figure details and remaining parameters as described in Fig. 4.

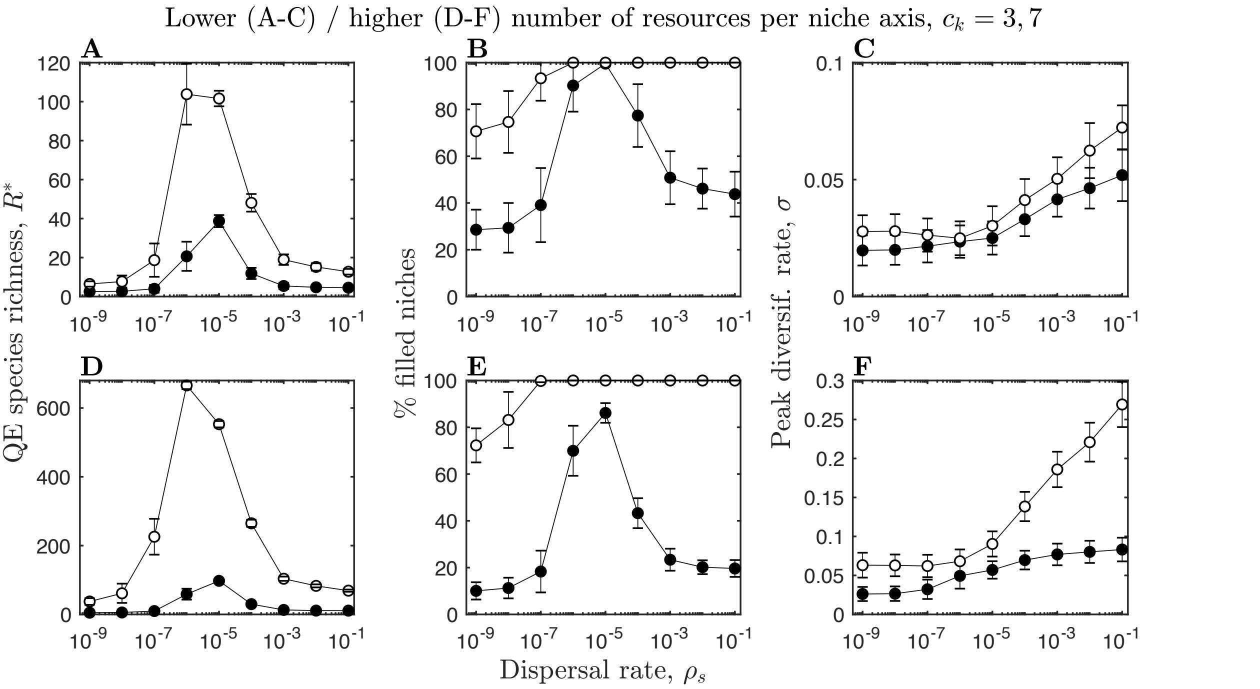


Figure S2 – Parameter sensitivity analysis for fewer (top row, $c_k = 3$) and more (bottom row, $c_k = 7$) resources per niche axis. Figure details and remaining parameters as described in Fig. 4.

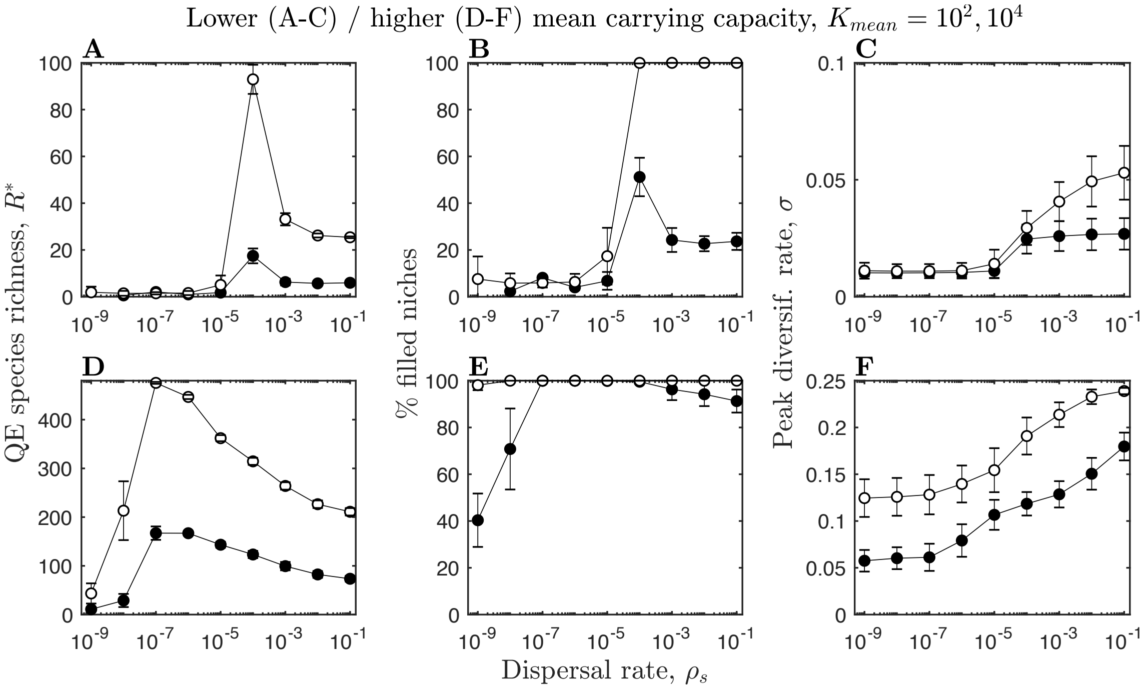


Figure S3 – Parameter sensitivity analysis for lower (top row, $K_{mean} = 10^2$) and higher (bottom row, $K_{mean} = 10^4$) mean carrying capacities. Figure details and remaining parameters as described in Fig. 4.

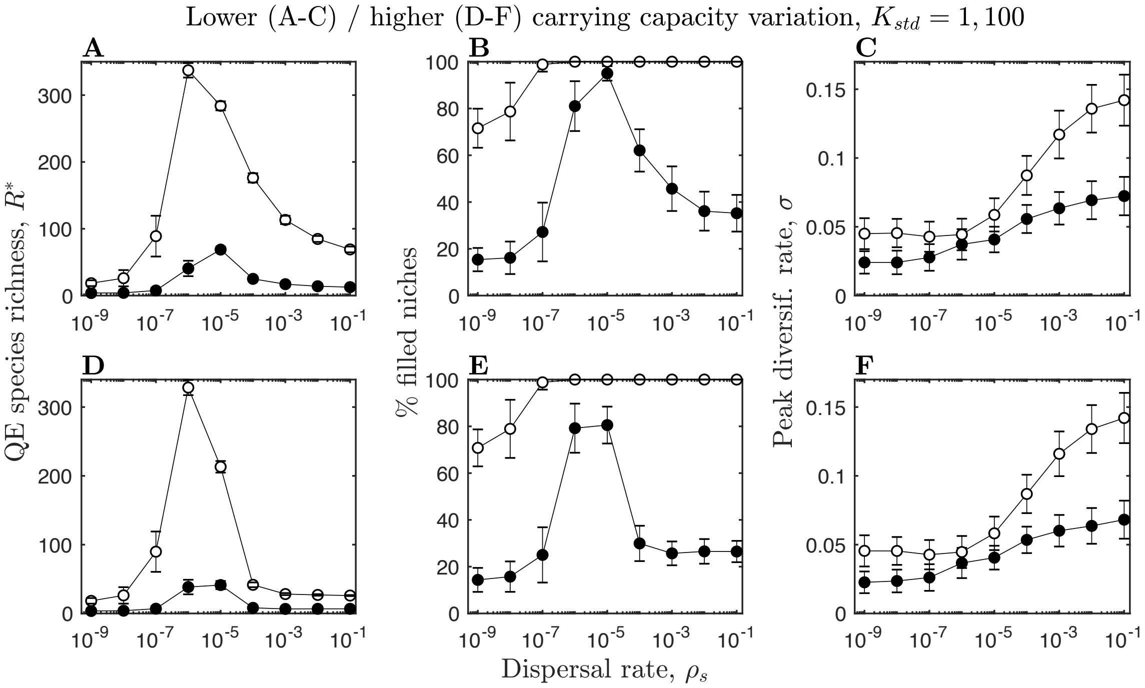


Figure S4 – Parameter sensitivity analysis for lower (top row, $K_{std} = 1$) and higher (bottom row, $K_{std} = 100$) standard deviations in carrying capacities. Figure details and remaining parameters as described in Fig. 4.

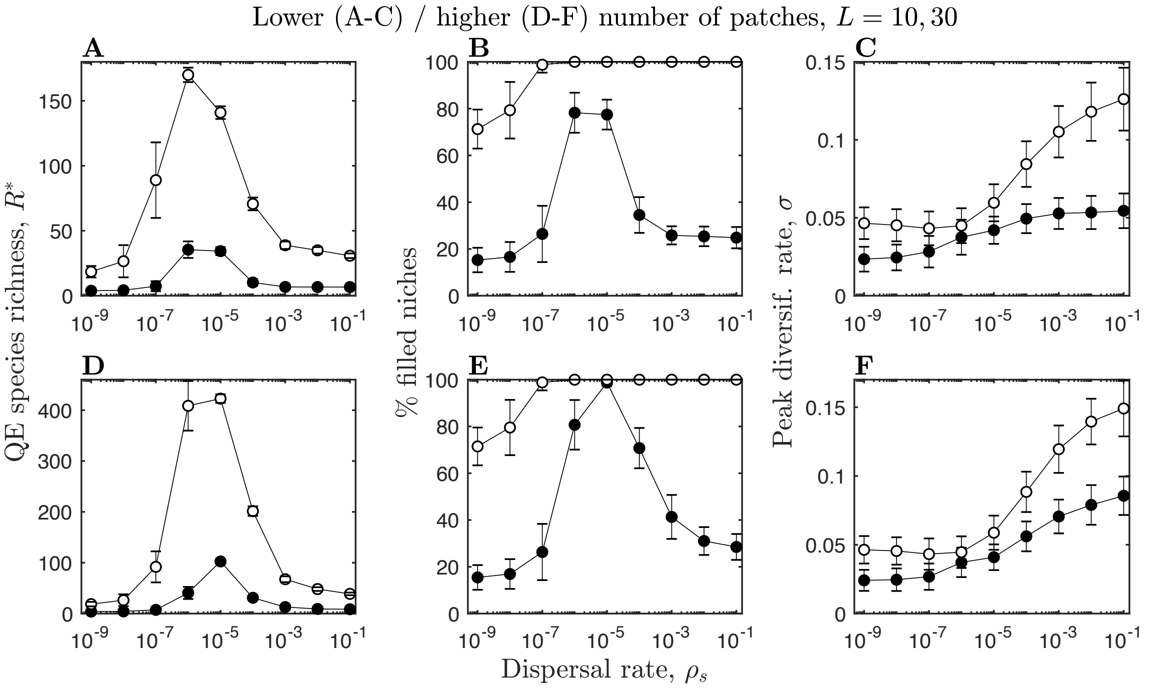


Figure S5 – Parameter sensitivity analysis for fewer (top row, $L = 10$) and more (bottom row, $L = 30$) patches. Figure details and remaining parameters as described in Fig. 4.

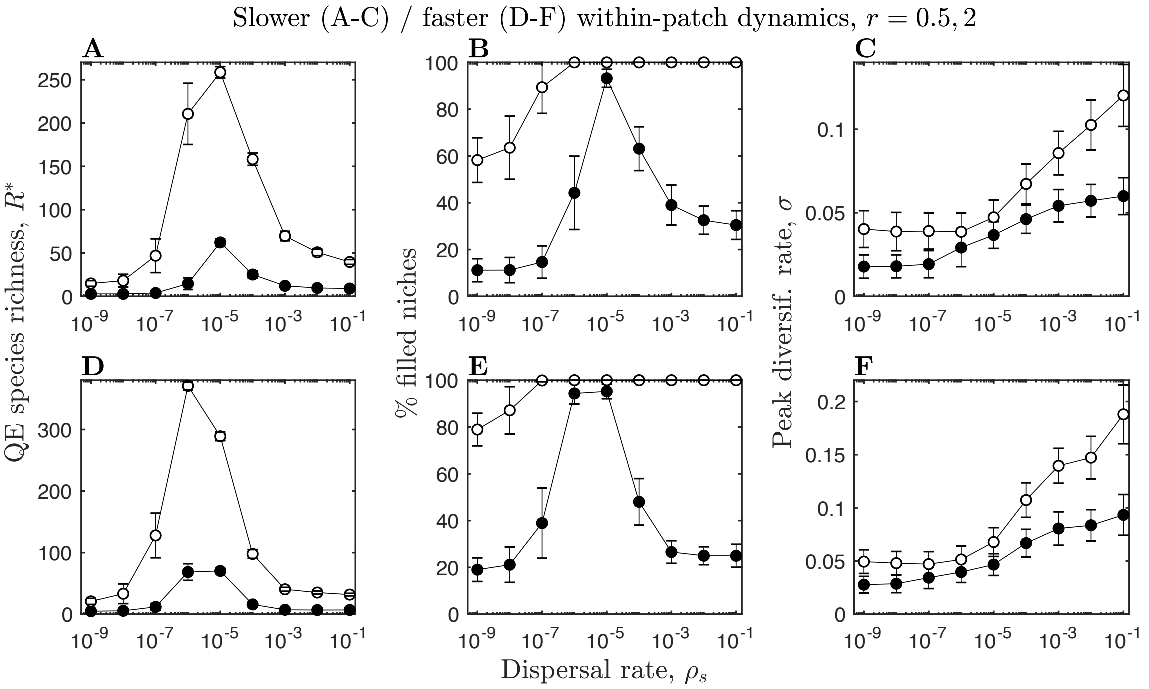


Figure S6 – Parameter sensitivity analysis for slower (top row, $r = 0.5$) and faster (bottom row, $r = 2$) within-patch dynamics. Figure details and remaining parameters as described in Fig. 4.

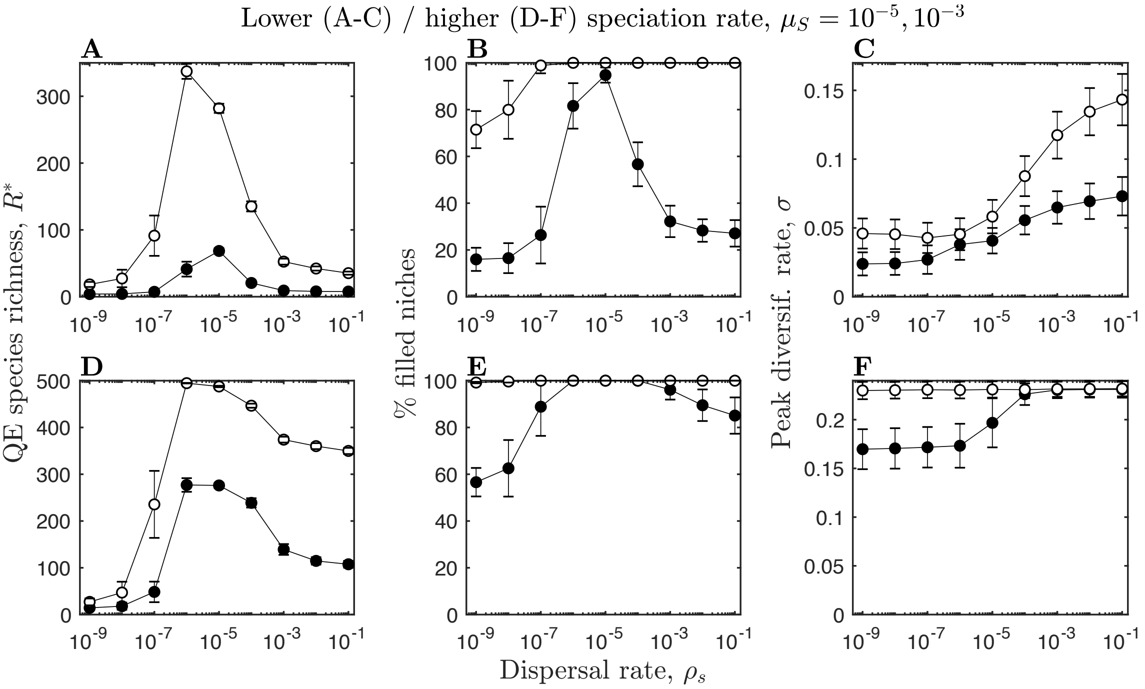


Figure S7 – Parameter sensitivity analysis for slower (top row, $\mu_S = 2 \times 10^{-4}$) and faster (bottom row, $\mu_S = 2 \times 10^{-3}$) speciation rates. Figure details and remaining parameters as described in Fig. 4.

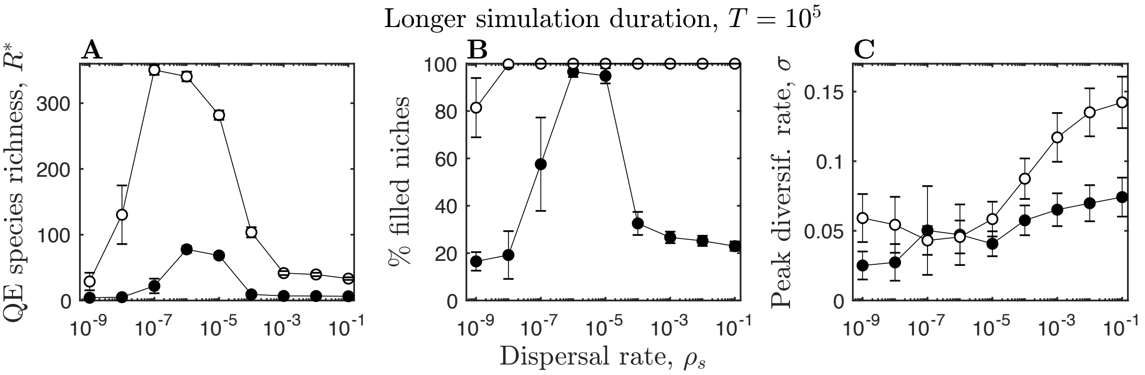


Figure S8 – Parameter sensitivity analysis for longer simulation durations ($T = 10^5$). Figure details and remaining parameters as described in Fig. 4.
