## Supplementary material for "An Inordinate Fondness for Species with Intermediate Dispersal Abilities": source code: competition_fast.c

```
/**************************************************************************
 * competition_fast.c
 * Fast competition function for file "dispersal_speciation.m'
 * output = competition_fast_v2(x,alpha,mk,L)
 * 
 * 20/05/2019
 * v1.0
 **************************************************************************/

#include <mex.h>
#include <math.h>
#include "matrix.h"

/*************************************
 * FUNCTION PROTOTYPES
 *************************************/
void** new2DArray(int row, int col, int typeSize);
void free2DArray(void **array, int row);

void mexFunction(int nlhs, mxArray *plhs[], int nrhs, const mxArray *prhs[])
{
    double *output, *x, *alpha, xsum, *parameter;
    int i, j, k, l, u, v, m1, m2, m3, m4, nofsites;
    const size_t *dims;
    
    /* Allocate inputs */    
    x= mxGetPr(prhs[0]);
    alpha= mxGetPr(prhs[1]);
    parameter= mxGetPr(prhs[2]);
    m1= (int)*parameter;
    parameter= mxGetPr(prhs[3]);
    nofsites= (int)*parameter;
    
    /* Get the dimensions in array */
    dims = mxGetDimensions(prhs[0]);
    m2 = m1*m1;
    m3 = m2*m1;
    m4 = m2*nofsites;
    
    /* Create output */
    if(nofsites==1){
        plhs[0] = mxCreateNumericArray(2, dims, mxDOUBLE_CLASS, mxREAL);
    }
    else{
        plhs[0] = mxCreateNumericArray(4, dims, mxDOUBLE_CLASS, mxREAL);
    }
    output = mxGetPr(plhs[0]);
    
    for(l=0;l<nofsites;l++){ /* Iterate through each patch */
        /* Sum over all identical niche species within this patch */
        for(j=0;j<m1;j++){
            for(i=0;i<m1;i++){
                output[i + j*m1 + l*m4] = 0;
                for(u=0;u<m1;u++){
                    for(v=0;v<m1;v++){
                        xsum = 0;
                        for(k=0;k<nofsites;k++) xsum += x[v + u*m1 + k*m2 + l*m4];
                        output[i + j*m1 + l*m4] += xsum*alpha[v + u*m1 + i*m2 + j*m3];
                    }
                }
                for(k=1;k<nofsites;k++) output[i + j*m1 + k*m2 + l*m4] = output[i + j*m1 + l*m4];
            }
        }
    }
    return;
}
```
