## Supplementary material for "An Inordinate Fondness for Species with Intermediate Dispersal Abilities": source code: dispersal_speciation.m

```
function [species_total,snapshots] = dispersal_speciation(max_time,r,Kmean,Kstd,rho,mu,epsilon,gk,ck,L,niche_type,sample_points)

%%%%%%%%%%%%%%%%%%%%%%%%%%%%%%%%%%%%%%%%%%%%%%%%%%%%%%%%%%%%%%%%%%%%%%%%%%%
% dispersal_speciation.m
%
% [species_total,snapshots] = dispersal_speciation(max_time,r,Kmean,Kstd,rho,mu,epsilon,gk,ck,L,niche_type,sample_points)
%
% SUMMARY:
% Source code for the stochastic model described in "An inordinate fondness
% for species with intermediate dispersal abilities."
% The model is implemented with 2 substitutable/non-substitutable niche
% traits, sensu Ashby et al. (2017) Competing species leave many potential
% niches unfilled. Nature Ecology & Evolution 1:1495-1501.
%
% DEPENDENCIES:
% - competition_fast.c: MEX file to be compiled with command 'mex
% competition_fast.c'.
% - poiss_rng.c: MEX file to be compiled with command 'mex poiss_rng.c'
%
% INPUTS:
% - max_time: duration of simulation
% - r: population dynamics rate (default = 1)
% - Kmean: mean carrying capacity (default = 1000)
% - Kstd: standard deviation in carrying capacity (default = 10)
% - rho: dispersal rate (default = 1e-5)
% - mu: speciation rate (default = 1e-4)
% - epsilon: extinction rate (default = 1e-3)
% - gk: parameter mediating niche breadth (default = 1.5)
% - ck: number of distinct traits on each niche axis (default = 5)
% - L: number of patches (default = 20)
% - niche_type: 0=non-substitutable resources, 1=substitutable resources (default = 0)
% - sample_points: time points to sample (default = 1:100:max_time)
%
% OUTPUTS:
% - species_total(t): global number of distinct species (combinations of niche
% - snapshots(i,j,k,t): the number of individuals with niche traits i and j
% in patch k at snapshot t
%
% The competition kernel, alpha(i,j,u,v) describes the strength of
% competition between species with niche traits (i,j) and (u,v).
%
% Let the difference between on niche axis 1 be E1 and on niche axis 2 be
% E2. If the niche axes are substitutable then:
%
% alpha(i,j,u,v) =exp(-gk*(E1+E2));
%
% and if they are non-substitutable then:
%
% alpha(i,j,u,v) = (exp(-gk*E1) + exp(-gk*E2))/2;
%
% where gk describes the strength of ecological competition and E1
% and E2 are the minimum distances between the two species' traits on each
% niche axis. Each niche axis is circular to remove boundary effects.

%%%%%%%%%%%%%%%%%%%%%%%%%%%%%%%%%%%%%%%%%%%%%%%%%%%%%%%%%%%%%%%%%%%%%%%%%%%

%%%%%%%%%%%%%%%%%%%%%%%%%%%%%%%%%%%%%%%%%%%%%%%%%%%%%%%%%%%%%%%%%%%%%%%%%%%
% Check dependencies
%%%%%%%%%%%%%%%%%%%%%%%%%%%%%%%%%%%%%%%%%%%%%%%%%%%%%%%%%%%%%%%%%%%%%%%%%%%
if(exist('poiss_rng','file')~=3)
    % No mex file exists, so try to compile
    disp('Attempting to compile poiss_rng.c ...')
    if(exist('poiss_rng.c','file')==0)
        disp('File poiss_rng.c not found.')
        poiss = str2func('poissrnd');
    else
        mex poiss_rng.c -silent
        disp('Compilation of poiss_rng.c complete.')
        poiss = str2func('poiss_rng');
    end
else
    poiss = str2func('poiss_rng');
end
if(exist('competition_fast','file')~=3)
    % No mex file exists, so try to compile
    disp('Attempting to compile competition_fast.c ...')
    if(exist('competition_fast.c','file')==0)
        disp('File competition_fast.c not found.')
        competition_fast_vers = 0;
    else
        mex competition_fast.c -silent
        disp('MEX compilation of competition_fast.c successful.')
        competition_fast_vers = 1;
    end
else
    competition_fast_vers = 1;
end

%%%%%%%%%%%%%%%%%%%%%%%%%%%%%%%%%%%%%%%%%%%%%%%%%%%%%%%%%%%%%%%%%%%%%%%%%%%
% Check inputs
%%%%%%%%%%%%%%%%%%%%%%%%%%%%%%%%%%%%%%%%%%%%%%%%%%%%%%%%%%%%%%%%%%%%%%%%%%%

if(nargin~=12)
    disp('Incorrect number of input arguments. Using defaults.');
    max_time = 1e4+1;
    r = 1;
    Kmean = 1000;
    Kstd = 10;
    rho = 1e-5;
    mu = 1e-4;
    epsilon = 1e-3;
    gk = 1.5;
    ck = 5;
    L = 20;
    niche_type = 0;
    sample_points = 1:100:max_time;
end
if(~isnumeric(max_time) || max_time<=0)
    disp('Parameter "Maxtime" must be positive. Using default: max_time = 1e4+1')
    max_time = 1e4+1;
end
max_time = ceil(max_time);

if(~isnumeric(r) || r<=0)
    disp('Parameter "r" must be positive. Using default: r = 1')
    r = 1;
end

if(~isnumeric(Kmean) || Kmean<=0)
    disp('Parameter "Kmean" must be positive. Using default: Kmean = 1000')
    Kmean = 1000;
end

if(~isnumeric(Kstd) || Kstd<0)
    disp('Parameter "Kstd" must be non-negative. Using default: Kstd = 10')
    Kstd = 10;
end

if(~isnumeric(rho) || rho<0)
    disp('Parameter "rho" must be non-negative. Using default: rho = 1e-5')
    rho = 1e-5;
end

if(~isnumeric(mu) || mu<=0)
    disp('Parameter "mu" must be positive. Using default: mu = 1e-4')
    mu = 1e-4;
end

if(~isnumeric(epsilon) || epsilon<0)
    disp('Parameter "epsilon" must be non-negative. Using default: epsilon = 1e-3')
    epsilon = 1e-3;
end

if(~isnumeric(gk) || gk<=0)
    disp('Parameter "gk" must be positive. Using default: gk = 1.5')
    gk = 1.5;
end

if(~isnumeric(ck) || ck<=0)
    disp('Parameter "ck" must be a positive integer. Using default: ck = 5')
    ck = 5;
end
ck = ceil(ck);

if(~isnumeric(L) || L<=0)
    disp('Parameter "L" must be a positive integer. Using default: L = 20')
    L = 20;
end
L = ceil(L);

if(~isnumeric(niche_type) || niche_type<0)
    disp('Parameter "niche_type" must be 0 or 1. Using default: niche_type = 0 (non-substitutable resources)')
    niche_type = 0;
end
niche_type = sign(niche_type);

if(~isnumeric(sample_points) || any(sample_points)<=0)
    disp('Parameter "sample_points" must be a non-negative array. Using default: sample_points = [0.1,1]*max_time')
    sample_points = 1:100:max_time;
end

%%%%%%%%%%%%%%%%%%%%%%%%%%%%%%%%%%%%%%%%%%%%%%%%%%%%%%%%%%%%%%%%%%%%%%%%%%%
% Create competition matrix
%%%%%%%%%%%%%%%%%%%%%%%%%%%%%%%%%%%%%%%%%%%%%%%%%%%%%%%%%%%%%%%%%%%%%%%%%%%

traits = (1:ck);
alpha = zeros(ck,ck,ck,ck);
for i1=1:ck
    for j1=1:ck
        E1 = min((traits(i1)-traits(j1))^2,min((ck+traits(i1)-traits(j1))^2,(traits(i1)-traits(j1)-ck)^2));
        for i2=1:ck
            for j2=1:ck
                if(i1==j1 && i2==j2)
                    alpha(i1,i2,j1,j2) = 1;
                else
                    E2 = min((traits(i2)-traits(j2))^2,min((ck+traits(i2)-traits(j2))^2,(traits(i2)-traits(j2)-ck)^2));
                    if(niche_type>0)
                        alpha(i1,i2,j1,j2) = exp(-gk*(E1+E2));
                    else
                        alpha(i1,i2,j1,j2) = (exp(-gk*E1) + exp(-gk*E2))/2;
                    end
                end
            end
        end
    end
end

%%%%%%%%%%%%%%%%%%%%%%%%%%%%%%%%%%%%%%%%%%%%%%%%%%%%%%%%%%%%%%%%%%%%%%%%%%%
% Create birth rate matrix & speciation mask
%%%%%%%%%%%%%%%%%%%%%%%%%%%%%%%%%%%%%%%%%%%%%%%%%%%%%%%%%%%%%%%%%%%%%%%%%%%
K = repmat(max(1e-30,Kmean + Kstd*randn(ck,ck,L)),[1,1,1,L]);
% r = repmat(permute(max(1e-30,rmean + rstd*randn(L,1)),[3,2,1]),[ck,ck,1,L]);
mask = zeros(ck,ck,L,L);
for i=1:L
    mask(:,:,i,i)=1;
end

%%%%%%%%%%%%%%%%%%%%%%%%%%%%%%%%%%%%%%%%%%%%%%%%%%%%%%%%%%%%%%%%%%%%%%%%%%%
% Set up population trackers and output
%%%%%%%%%%%%%%%%%%%%%%%%%%%%%%%%%%%%%%%%%%%%%%%%%%%%%%%%%%%%%%%%%%%%%%%%%%%

% Population tracker
x = zeros(ck,ck,L,L);
x(ceil(ck/2),ceil(ck/2),1,1) = Kmean;
snapshots = zeros(ck,ck,L,L,length(sample_points));

% Output
species_total = zeros(length(sample_points),1);
snapshotcount = 1;

%%%%%%%%%%%%%%%%%%%%%%%%%%%%%%%%%%%%%%%%%%%%%%%%%%%%%%%%%%%%%%%%%%%%%%%%%%%
% Carry out the simulation
%%%%%%%%%%%%%%%%%%%%%%%%%%%%%%%%%%%%%%%%%%%%%%%%%%%%%%%%%%%%%%%%%%%%%%%%%%%

for t=1:max_time
    
    % Global extinction
    if(all(x==0))
        break
    end
    
    %%% 1) Population dynamics %%%
    
    % Competition within patches
    if(competition_fast_vers>0 && L>1)
        competition = competition_fast(x,alpha,ck,L);
    else
        % Non-MEX code
        competition = zeros(ck,ck,L,L);
        for k=1:L
            competition(:,:,:,k) = repmat(permute(sum(sum(alpha.*repmat(sum(x(:,:,:,k),3),[1,1,ck,ck]),2),1),[3,4,1,2]),[1,1,L,1]);
        end
    end
    
    % Calculate net change in population size
    x = poiss(x.*(1+r*(1-competition./K)));
    
    %%% 2) Speciation %%%
    x0 = sum(x,3);
    speciations_out1 = poiss(0.25*mu*x0);
    speciations_out2 = poiss(0.25*mu*x0);
    speciations_out3 = poiss(0.25*mu*x0);
    speciations_out4 = poiss(0.25*mu*x0);
    temp = speciations_out1([2:end,1],:,:,:) + speciations_out2([end,1:(end-1)],:,:,:) + speciations_out3(:,[2:end,1],:,:) + speciations_out4(:,[end,1:(end-1)],:,:);
    speciations = mask.*repmat(temp,[1,1,L,1]); % Patch identifiers for where mutants arise
    
    %%% 3) Dispersal %%%
    dispersal_out1 = min(x,poiss(0.5*rho*x));
    dispersal_out2 = min(x,poiss(0.5*rho*x));
    dispersal_out = dispersal_out1 + dispersal_out2;
    
    diff1 = max(0,(dispersal_out-x));
    temp1 = round(diff1.*(dispersal_out1./max(1,dispersal_out)));
    temp2 = diff1-temp1;
    
    dispersal_out1 = dispersal_out1 - temp1;
    dispersal_out2 = dispersal_out2 - temp2;
    dispersal_out = dispersal_out1 + dispersal_out2;
    dispersal_in = dispersal_out1(:,:,:,[2:end,1]) + dispersal_out2(:,:,:,[end,1:(end-1)]);
    
    % Update population
    x = max(0,x + speciations + dispersal_in - dispersal_out);
    
    %%% 4) Extinctions %%%
    if(epsilon>0)
        alive = find(x);
        extinctions = alive(rand(length(alive),1)<epsilon);
        x(extinctions) = 0;
    end
    
    %%% 5) Update sample_points of species total %%%
    if(snapshotcount<=length(sample_points))
        if((t+1)>=sample_points(snapshotcount))
            snapshots(:,:,:,:,snapshotcount) = x;
            species_total(snapshotcount) = length(find(sum(x,4)>0));
            snapshotcount = snapshotcount + 1;
        end
    end
end
```
