## Supplementary material for "An Inordinate Fondness for Species with Intermediate Dispersal Abilities": source code: poiss_rng.c

```
/**************************************************************************
 * poiss_rng.c
 * Poisson random number generator
 * Given an input array y = [y1,y2,...yn], this function generates an
 * equally sized array of random numbers from poisson distributions with
 * mean values given by y1,y2,...,yn. If an input value is large, use the 
 * method proposed by Ahrens and Dieter in Knuth, Volume 2, 1998 edition.
 * 
 * 24/07/2018
 * v1.0
 *************************************************************************/
#include "mex.h"
#include "math.h"
#define PI 3.141592653589793

/* Don't change these values, used to generate pseudorandom numbers */
static unsigned int m_w = 521288629;
static unsigned int m_z = 362436069;

double binomial_rng(long, double);
double gamma_rng(double);
double normal_rng(void);
double poiss_small(double);
double poiss_large(double);
double uniform_rng(void);

/* Mex gateway function */
void mexFunction( int nlhs, mxArray *plhs[],
        int nrhs, const mxArray *prhs[])
{
    double *y,*z,*parameter;
    int i, newseed;
    size_t ndim;
    const size_t *dims;
    size_t elements;
    
    /* Input array */
    y = mxGetPr(prhs[0]);
    
    /* Get the dimensions in array */
    ndim=mxGetNumberOfDimensions(prhs[0]);
    dims = mxGetDimensions(prhs[0]);
    
    /* Get the number of elements in the input argument */
    elements=mxGetNumberOfElements(prhs[0]);
    
    /* Create output array */
    plhs[0] = mxCreateNumericArray(ndim, dims, mxDOUBLE_CLASS, mxREAL);
    z = mxGetPr(plhs[0]);
    
    /* Main loop */
    z[0] = 0;
    for(i=0;i<elements;i++){
        if(y[i]<=0){
            z[i] = 0;
        }
        else if(y[i]<15){
            z[i] = poiss_small(y[i]);
        }
        else{
            z[i] = poiss_large(y[i]);
        }
    }
}

double binomial_rng(long n, double p)
/**************************************************************************
 * Returns a normally distributed integer between 0 and n inclusive
 *************************************************************************/
{
    long i;
    double x = 0;
    
    for (i = 0; i < n; i++) x += ((uniform_rng() < (1.0 - p)) ? 0 : 1);
    return (x);
}

double gamma_rng(double shape)
/**************************************************************************
 * Returns a gamma distributed positive real number
 * Based on Marsaglia G and Tsang W.W. (2000) A Simple Method for Generating
 * Gamma Variables. ACM Transactions on Mathematical Software, 26:363-372
 *************************************************************************/
{
    double d, c, x, xsquared, v, u;
    double scale = 1;
    
    if (shape >= 1.0){
        d = shape - 1.0/3.0;
        c = 1.0/sqrt(9.0*d);
        for (;;){
            do{
                x = normal_rng();
                v = 1.0 + c*x;
            }
            while (v <= 0.0);
            v = v*v*v;
            u = uniform_rng();
            xsquared = x*x;
            if (u < 1.0 -.0331*xsquared*xsquared || log(u) < 0.5*xsquared + d*(1.0 - v + log(v))){
                return scale*d*v;
            }
            else {
                double g = gamma_rng(shape+1.0);
                double w = uniform_rng();
                return scale*g*pow(w, 1.0/shape);
            }
        }
    }
    return false;
}

double normal_rng(void)
/**************************************************************************
 * Returns a normally distributed real number, using the Box-Muller
 * algorithm
 *************************************************************************/
{
    double u1 = uniform_rng();
    double u2 = uniform_rng();
    double r = sqrt( -2.0*log(u1) );
    double theta = 2.0*PI*u2;
    return r*sin(theta);
}

double poiss_large(double y1)
/**************************************************************************
 * Use this function for large input values
 *************************************************************************/
{
    double z1=0;
    double m,x;
    
    m = floor(y1*7/8);
    x = gamma_rng(m);
    if(x<y1) {
        if((y1-x)<15){
            z1 = m + poiss_small(y1-x);
        }
        else {
            z1 = m + poiss_large(y1-x);
        }
    }
    else {
        z1 = binomial_rng((long)m-1, y1/x);
    }
    return z1;
}

double poiss_small(double y1)
/**************************************************************************
 * Use this function for small input values
 *************************************************************************/
{
    double z1=-1;
    double t = 0;
    
    while (t <= y1) {
        t += (-log(1.0 - uniform_rng()));
        z1++;
    }
    return z1;
}

double uniform_rng(void)
/**************************************************************************
 * Returns a uniformly distributed real number
 *************************************************************************/
{
    unsigned int u;
    m_z = 36969 * (m_z & 65535) + (m_z >> 16);
    m_w = 18000 * (m_w & 65535) + (m_w >> 16);
    u = (m_z << 16) + m_w;
    /* Scale between 0 and 1: 2.328306435454494e-10 = 1/(2^32 + 2) */
    return (u + 1.0) * 2.328306435454494e-10;
}
```
